# Thermal proteome profiling identifies new drug targets in Plasmodium falciparum parasites

**DOI:** 10.64898/2026.01.30.702724

**Authors:** Samuel Pazicky, Jerzy M. Dziekan, Seth Tjia, Selina Bopp, Dyann Wirth, Zbynek Bozdech

**Author notes:** Corresponding author: Zbynek Bozdech.

## Abstract

Spreading resistance to clinically used antimalarial drugs increases the need for identification of new drug targets. Here, we screened 25 antimalarials with known and unknown mode of action to validate old and find new drug targets in *P. falciparum*. Combining experimental approach by proteome-wide cellular thermal shift assay and computational filtering by molecular docking, we found the drugs to bind to previously known drug targets, validated ACS10 as the target of MMV665915 and discovered new drug targets. Furthermore, we updated the experimental and analytical MS-CETSA pipeline with the inclusion of membrane solubilization step, validated the targets of atovaquone and cipargamin on cell lysates and in intact cells and mapped the parasite response to these antimalarials, identifying monocarboxylate transporter MCP2 as atovaquone transporter.

## Introduction

Counting almost 250 million cases and over 600 thousand deaths in 2023 alone, malaria remains a global threat to public health^1^. Since the break of the millennium, artemisinin combination therapy (ACT) has been instrumental in reducing malaria-related deaths worldwide^2^. Unfortunately, the efficacy of ACTs has declined over the past years as the malaria parasites, *Plasmodium falciparum*, began to developed resistance to artemisinin and consequently to most of the combination therapy partner drugs^3^. Emerging in Southeast Asia in 2009, artemisinin resistance underlines the progressively increasing frequencies of ACT failures withing the region^4–6^. The recent emergence of artemisinin resistance of *P. falciparum* in Sub-Saharan Africa is thus deeply disconcerting^7^, emphasizing the need for replacement chemotherapeutics. Discovery of new drugs and identification of new drug targets are thus among the highest priorities for all global control and elimination of malaria programs^8^.

Historically, most of the antimalarial drugs were identified in large-scale phenotypic screens, being agnostic to their mechanisms of action (MoA)^9^. Direct drug targets for a small number of antimalarial drugs were, however, ultimately discovered such as dihydrofolate reductase (DHFR, targeted by proguanil and pyrimethamine)^10^, dihydropteroate synthase (DHPS, targeted by sulfadoxine)^11^, and mitochondrial bc1 complex (targeted by atovaquone)^12^. Most of the current antimalarial drug discovery efforts also start with high throughput phenotypic screens (HTS) but much more emphasis is being put on the subsequent characterizations of the MoA and drug targets^13^. Such information is crucial for further compound optimizations to maximize its potency against the parasite growth *per se* but also to streamline the *in-patient* evaluations before possible clinical trials^14^. *In vitro* evolution and whole genome analysis (IVIEWGA) became the gold standard method of deciphering the new antimalarial compound’s MoAs with great success^13^. IVIEWGA was used to identify a considerable number of antimalarial drug targets over the last decade including PfATP4 (targeted by cipargamin^15,16^); elongation factor 2 (Cabamiquine, M5717)^17^; cyclin-dependent–like (CLK3) protein kinase (TCMDC-13505)^18^; Acyl-CoA synthetases, ACS10/11, (GSK701)^19^ and many others (reviewed Flannery *et al*^8^).

IVIEWGA has, however, several challenges that can in some cases obscure genuine target discoveries. These include rising mutations in factors of resistance mechanisms^20 21^, and/or fitness alleviating factors^13^ on one side and inability to derive any resistant parasites with putative genetic variations on the other^13,22^. These challenges can be overcome by a recently developed alternative method known as thermal protein profiling (TPP) or cellular thermal shift assay (CETSA), which we have previously introduced for drug target identifications in *P. falciparum*^23^. CETSA is based on the biophysical phenomenon that by binding to a drug, the target protein becomes more resistant to aggregation by thermal denaturation^24,25^. Coupled with mass spectroscopy-based quantitative proteomics, CETSA can determine protein stabilities proteome-wide and as such allows discoveries of direct protein-drug biding *de novo*. Over the last five years, CETSA has proven to be a highly effective high-throughput method for drug target identification in various systems such as cancer^26^ but also across a large spectrum of infectious pathogens^27–30^. Optimized for malaria parasites (*P. falciparum*), CETSA can detect protein-drug engagement either in total protein lysates (*lysate* CETSA) or in intact cells within its erythrocyte host (*in-cell* CETSA), cross-referring of both can generate high confidence predictions for drug target interactions^23^. This was initially observed for clinical antimalarial drugs such as pyrimethamine-DHFR-TS^27^ but also an experimental compound MMV00848-falcilysin interactions^31^. Over the last five years, CETSA proved to be useful in identifying direct protein targets in *P. falciparum* for several experimental compounds including the eukaryotic translation initiation factor 3 (eIF3I) inhibited by quinazoline-quinoline bi-substrates^32^; PfFKPB35 and other interacting partners of FK506^33^; purine nucleoside phosphorylase (PfPNP) interacting with MMV00848, and multiple targets of chloroquine^34^. CETSA was also instrumental in uncovering a host factor, PRDX6, a lipid-peroxidation repair enzyme with phospholipase A2 (PLA2), that can block malaria parasite growth, being inhibited by darapladib^35^.

Over the last decade, CETSA-based methodologies have emerged as increasingly informative approaches to drug target identification, relevant to current but also future antimalarial drug discoveries^13,23,36^. As such, CETSA-based methodologies warrant further development and optimization to enhance their informality for future antimalarial and other antimicrobial drug discovery efforts. Here, we applied CETSA to protein target identification for a panel of antimalarial compounds to provide a broader, comprehensive overview of the efficiency of these technologies for protein target discovery in the key malaria pathogen *P. falciparum*. In the process, we developed several key methodological improvements and demonstrated their utility to substantially improve such analyses in the future.

## Results

### Identification of *P. falciparum* putative drug targets using an improved bioinformatics pipeline (ITDRMS)

First, we applied the previously developed isothermal dose response (ITDR) format of CETSA to identify proteins exhibiting significant thermal stability shifts induced by a set of 25 antimalarial experimental drug candidates (Supplementary Tables 1 and 2). In ITDR format, a range of compound concentration is used to measure dose-dependent changes in protein solubility at constant temperatures. Specifically, we performed ITDR CETSA with protein lysates extracted from in vitro cultured *P. falciparum* at the late trophozoite stage that presumably reflect direct drug binding, as these are not affected by downstream effects that can occur in intact cells. Across all measurements, we detected for 3,702 parasite proteins that could be fitted with dose-response curves, of which 1,617 were fitted in all 25 datasets and another 657 in at least 20 datasets (Extended Data Fig. 1a). Subsequently, we developed a new in-house R package, *ITDRMS,* that evaluates each protein’s dose-dependent thermal profile using custom-designed scaling and fitting of each thermal profile and subsequent statistical evaluation of their significance (see Materials and Methods). *ITDRMS* is based on a statistical description of the fitted curves by confidence intervals for hit calling as opposed to previously applied *mineCETSA*^23^, *which* uses rigid cutoff parameters for R^2^ of the curve fitting (default 0.8). Comparing the *ITDMRS* and *mineCETSA* applied to the CETSA measurements for the 25 antimalarial drugs (Extended Data Fig. 1b), we can show that *ITDMRS* rejects putative false-positive thermal shifts driven by single outliers in the lowest or highest drug dose and removes ambiguous hits that display simultaneous stabilization and destabilization at different temperatures (Extended Data Fig. 1c). On the other hand, *ITDRMS* can identify some putative drug targets that were excluded by strict cutoffs in *mineCETSA* package (Extended Data Fig. 1d).

Applying the newly developed *ITDRMS* pipeline to the aforementioned dataset, we identified 99 proteins (CI<0.05, response>1) stabilized by at least one of the 25 compounds studied (Fig. 1a, Supplementary Table 3). While most compounds stabilized only 1-4 proteins (out of 21 compounds), three compounds (MMV665806, MMV666070, and MMV007685) stabilized 13, 21, and 45 proteins, respectively, and MMV665875 yielded no significant thermal shifts. With the exception of these three multitarget compounds, we observed minimal overlap among the stabilized proteins induced by individual compounds, indicating the specificity and selectivity of each compound (Fig. 1a). This is further supported by the fact that our *ITDRMS* analysis could replicate previously validated ligand-protein binding for several well-established drug targets. These include the cladosporin target, lysyl tRNA synthetase (Fig. 1b)^37^ and the MMV000848 target, PfPNP (Extended Data Fig. 2a)^27^. Interestingly, two additional compounds (MMV666070 and MMV007695) also significantly stabilized PfPNP, suggesting somewhat more promiscuous binding capacity of its active site (Extended Data Fig. 2a). Another example represents *Plasmodium* protease falcilysin, whose previously reported binding to MMV000848 and MMV665806^31^ was captured by *ITDRMS* analysis (Extended Data Fig. 2b).

**Figure 1.**
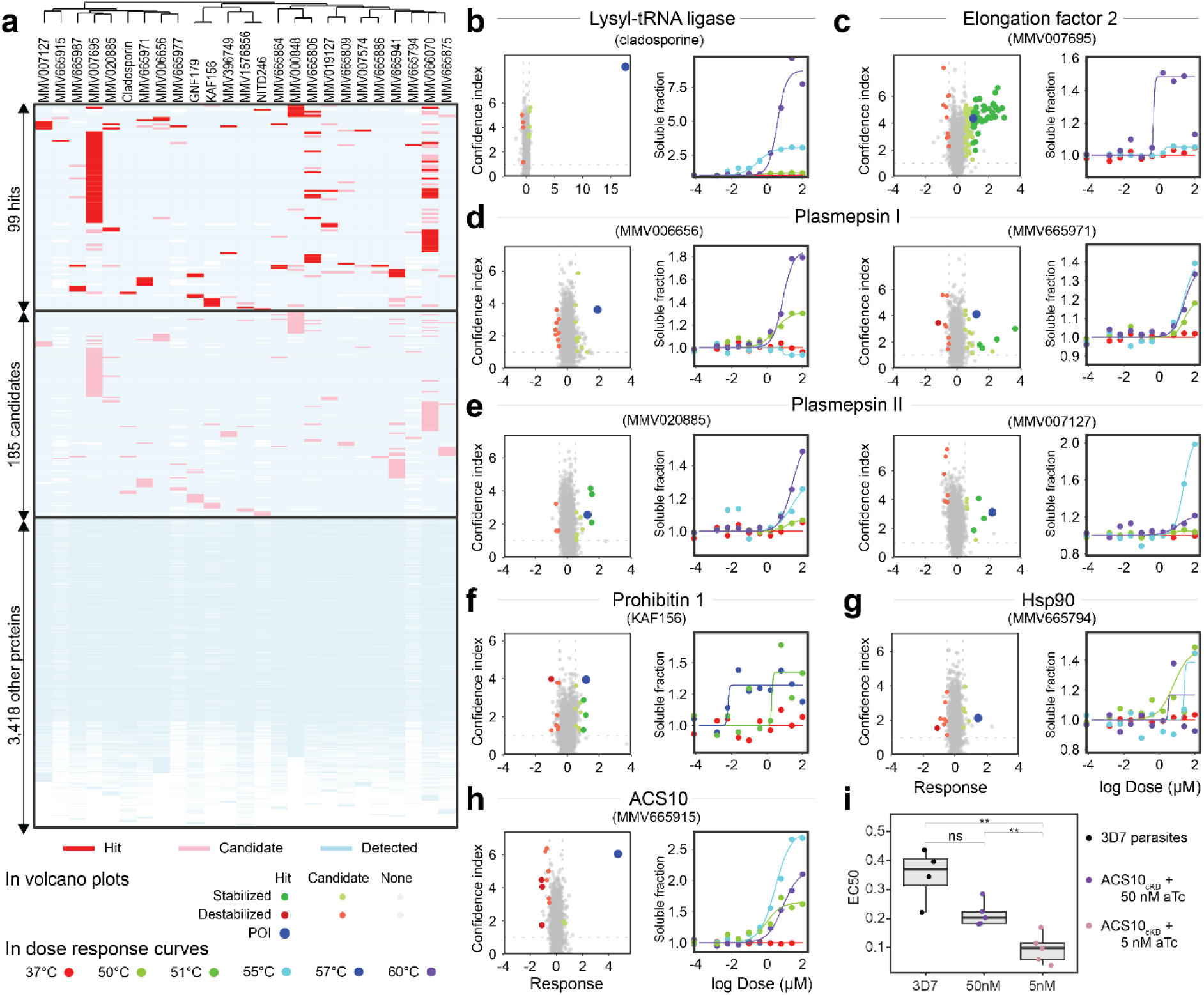
MS-CETSA screen of 25 antimalarials. **a** Heatmap showing 99 protein hits (red), 185 candidates (pink) and 3566 non-interacting proteins (light blue). Undetected proteins are white, the compounds are clustered based on the similarity of their molecular fingerprints. **b-h** Volcano plots (left) and dose-response curves (right) of hit-compound pairs. In volcano plots, green are stabilized hits, light green are stabilized compounds, red are destabilized hits and pink are destabilized candidates. Other proteins are grey, and the proteins of interest are blue. The plots show dose-response curves of the proteins of interest at either 37 °C (control temperature, red), 50°C (light green), 51°C (green), 55°C (cyan), 57°C (blue) or 60°C (purple). **i** Significant decrease of EC50 in low ACS10 condition (5 µM aTc) compared to normal level ACS10 (50 µM aTc) and wild-type parasites (Student’s t-test) validates ACS10 as MMV665915 target).

In addition to these well-established protein-ligand pairs, we identified several proteins whose potential as antimalarial drug targets has been or is currently being evaluated, albeit here bound by alternative chemical structures. First, we identified elongation factor 2 (PfEF2, PF3D7_1451100), thermally stabilized by MMV007695 (Fig. 1c). PfEF2 is believed to be the target of M5717, a quinoline-4-carboxamide derivative active in liver and blood stages, which is currently undergoing clinical trials^17,38^. Second, two proteases involved in hemoglobin digestion, plasmepsins I and II, were stabilized each by two compounds, including MMV665971 and MMV006656 (Fig. 1d), and MMV020885 and MMV007127 (Fig. 1e). Both pairs of compounds display structural similarities (Supplementary Table 2) that may explain the overlaps between their stabilization profiles. Third, we identified prohibitin 1 (PF3D7_0829200) as one of the strongest interacting targets of KAF156 (Fig. 1f). Inhibition of prohibitin 1 has previously been shown to abrogate parasite growth^39^. KAF156 (better known as ganaplacide) has been evaluated in Phase II clinical trials, including in combination with lumefantrine, showing rapid parasite clearance in patients with uncomplicated malaria, but its MoA is currently unknown^40^. Fourth, the most stabilized protein by MMV665794 was heat shock protein PfHsp90 (PF3D7_0708400, Fig. 1g), which was also among the hits of MMV007695 (Extended Data Fig. 2c). PfHsp90 inhibitors have been previously identified from a screen of pharmaceuticals and FDA-approved drugs, and they were found to inhibit the parasite growth^41–43^. Finally, we found a previously characterized antimalarial drug target, acyl-CoA synthetase ACS10 (PF3D7_0525100)^19^, which here was the only protein stabilized by MMV665915 (Fig. 1h) and one of many proteins stabilized by MMV007695 (Extended Data Fig. 2d). To validate ACS10 as MMV665915 target, we used the previously generated PfACS10 conditional knockdown parasites (ACS10_cKD_) that express ACS10 only in the presence of anhydrotetracycline (aTc)^19^. In agreement with the screen results, MMV665915 is more potent when the abundance of ACS10 is low (5 µM aTc, EC50=0.09 µM) compared to normal abundance (50 µM aTc, EC50=0.20 µM) and wild type parasites (EC50=0.37 µM) (Fig. 1i). Taken together, in addition to the previously known ligand-protein binding (above), our CETSA screen identified several protein factors whose targeting has a high parasiticidal potential.

Overall, our 25-compound CETSA screen identified as many as 99 proteins, most of which were not previously known to interact with any compounds with *in vitro* antimalarial activities. Here we wish to highlight six of these that, similarly to the aforementioned protein targets, were exclusive to a corresponding compound or the thermal stabilization reached the highest statistical confidence. These included diphthine methyl ester synthase DPH5 (*PF3D7_1009000*), which was the only *Plasmodium* protein stabilized by the MMV665977, and the “top hit” for MMV007127 and MMV396749 (Fig. 2a). Similarly, mitochondrial import inner membrane translocase TIM50 (*PF3D7_0726900*) was the only protein stabilized by MMV665886, and a highly confident hit for MMV665864 and MMV665987 (Fig. 2b). Nucleoside transporter ENT1 (*PF3D7_1347200*), whose disruption in *P. berghei* was shown to suppress cerebral malaria^44^, was significantly stabilized by MMV666070 and MMV020885 (Fig. 2c). Finally, hydroxyethylthiazole kinase (*PF3D7_1239600*), nucleotidyl transferase (*PF3D7_1332400*), and ATP-dependent RNA helicase DDX41 (*PF3D7_0527900*), were the top hits of GNF179, MMV007574 and MMV665809, respectively (Fig. 2d-f). In line with all the presented evidence, it is feasible to suggest that the thermal shifts of these novel protein factors represent genuine binding of the studied compounds and that this (presumably) inhibitory binding mediates and/or contributes to their parasiticidal mode of action (MoA).

**Figure 2.**
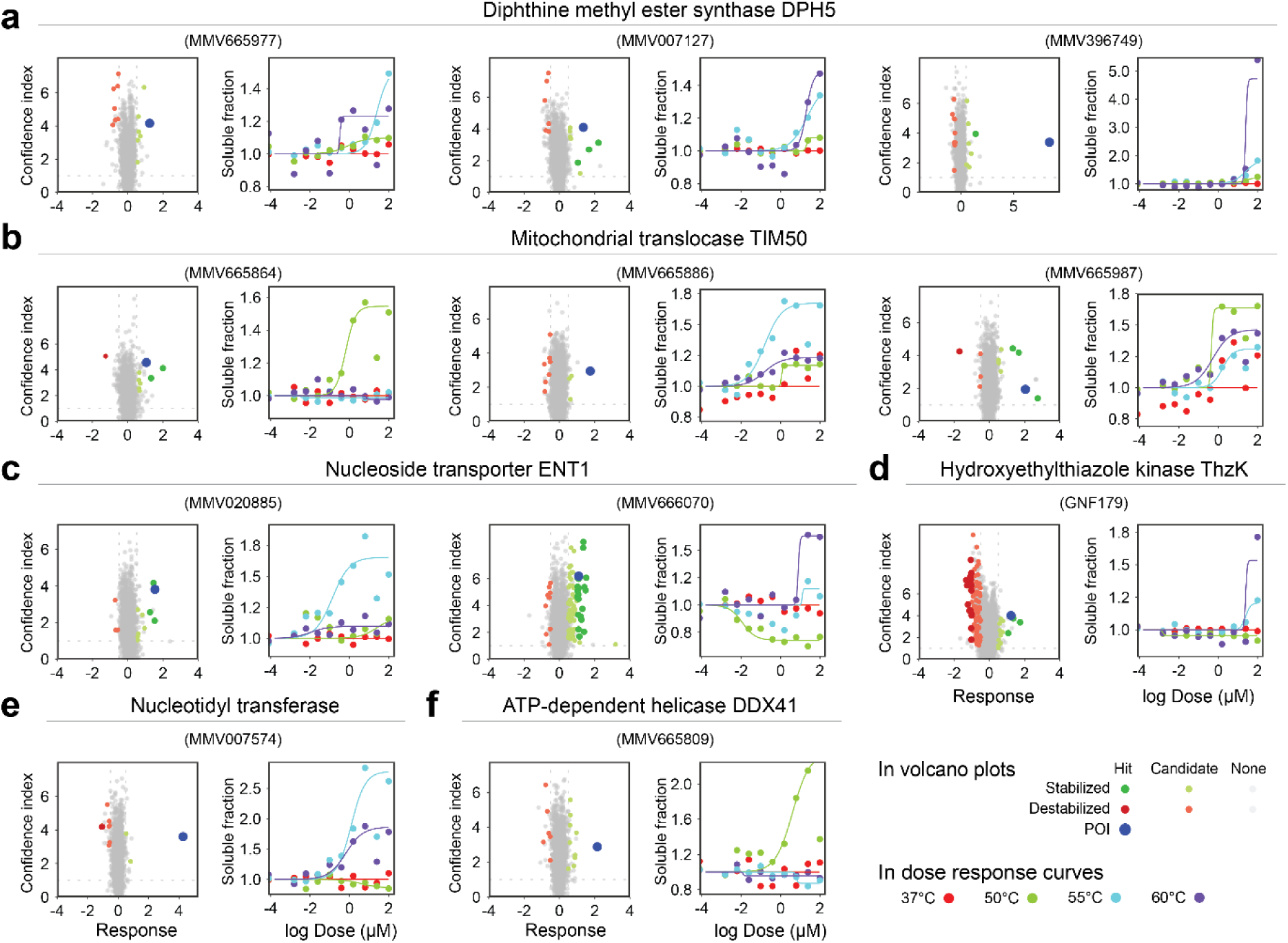
New high-confident hits identified by MS-CETSA screen. Volcano plots (left) and dose-response curves (right) of hit-compound pairs. In volcano plots, green are stabilized hits, light green are stabilized compounds, red are destabilized hits and pink are destabilized candidates. Other proteins are grey, and the protein of interest is blue. The right plots show dose-response curves of the proteins of interest at either 37 °C (control temperature, red), 50°C (light green), 55°C (cyan), or 60°C (purple).

### Molecular docking predicts competitive inhibition of target enzymes

To support the suggestive evidence of the protein-compound binding identified by the CETSA screen, we employed *in silico* molecular docking as an orthogonal method to prioritize the selection of drug candidates. As an initial we considered the highly confident 99 *P. falciparum* and an additional 185 protein-compound binding pairs whose compound-induced thermal shift confidences were above somewhat relaxed thresholds (CI < 0.1; response > 0.5) (Fig. 1a). For the *in silico* structural analysis, we first selected 92 proteins that bind multiple compounds and, after filtering, considered 35 proteins that were presumably monomeric, soluble, cytoplasmic proteins with experimentally determined structures for their putative homologs (similarity > 50%) with bound ligands (Fig. 3a, Extended Data Fig. 3a). For each protein-compound interaction, the docking pipeline generated up to 10 binding conformations (poses) with the lowest calculated binding energies. In our analysis, the number of docking poses proximal to the experimentally confirmed binding pockets appeared highly predictive of true binding compared to the calculated binding affinities *per se* (Extended Data Fig. 3b). For example, the mean binding affinity of cladosporin docked in its target, lysyl-tRNA ligase, was only -7.0 kcal/mol, but 7 out of ten poses docked in the close vicinity of the binding pocket (Extended Data Fig. 3c, Supplementary Table 4). Similarly, MMV665806 and MMV000848 docked in one of the two mefloquine binding pockets of falcilysin, albeit with relatively low calculated binding energies (Extended Data Fig. 3d, Supplementary Table 4). Hence, we chose the targets with 5 or more poses in a binding pocket as high-confidence hits from which we could prioritize 13 putative drug targets (Supplementary Table 4 and Supplementary Data 1). These include diphthine methyl ester synthase DPH5 (Fig. 2a) and nucleoside transporter ENT1 (Fig. 2c), for which the in silico docking confirmed the putative binding of their respective compounds within their active sites (Fig. 3bc). Another example is peptide deformylase (PDF, PF3D7_0907900) putative target of MMV665915 and MMV665977 both of which were predicted to share a binding site with the previously identified PDF inhibitor (Fig. 3d). The PDF homologues have been extensively studied as antibacterial targets, although their existing PDF inhibitors appear to bind to zinc metalloprotease FTSH1 in *Plasmodium*^45^. Likewise, the interactions between phosphoethanolamine methyltransferase (PF3D7_1343000) and both MMV666070 and MMV665806 identified by our CETSA screen were predicted to occur in one of the two binding sites previously predicted for inhibitory binding by amodiaquine ^46^ suggesting their inhibitory potential (Fig. 3e).

**Figure 3.**
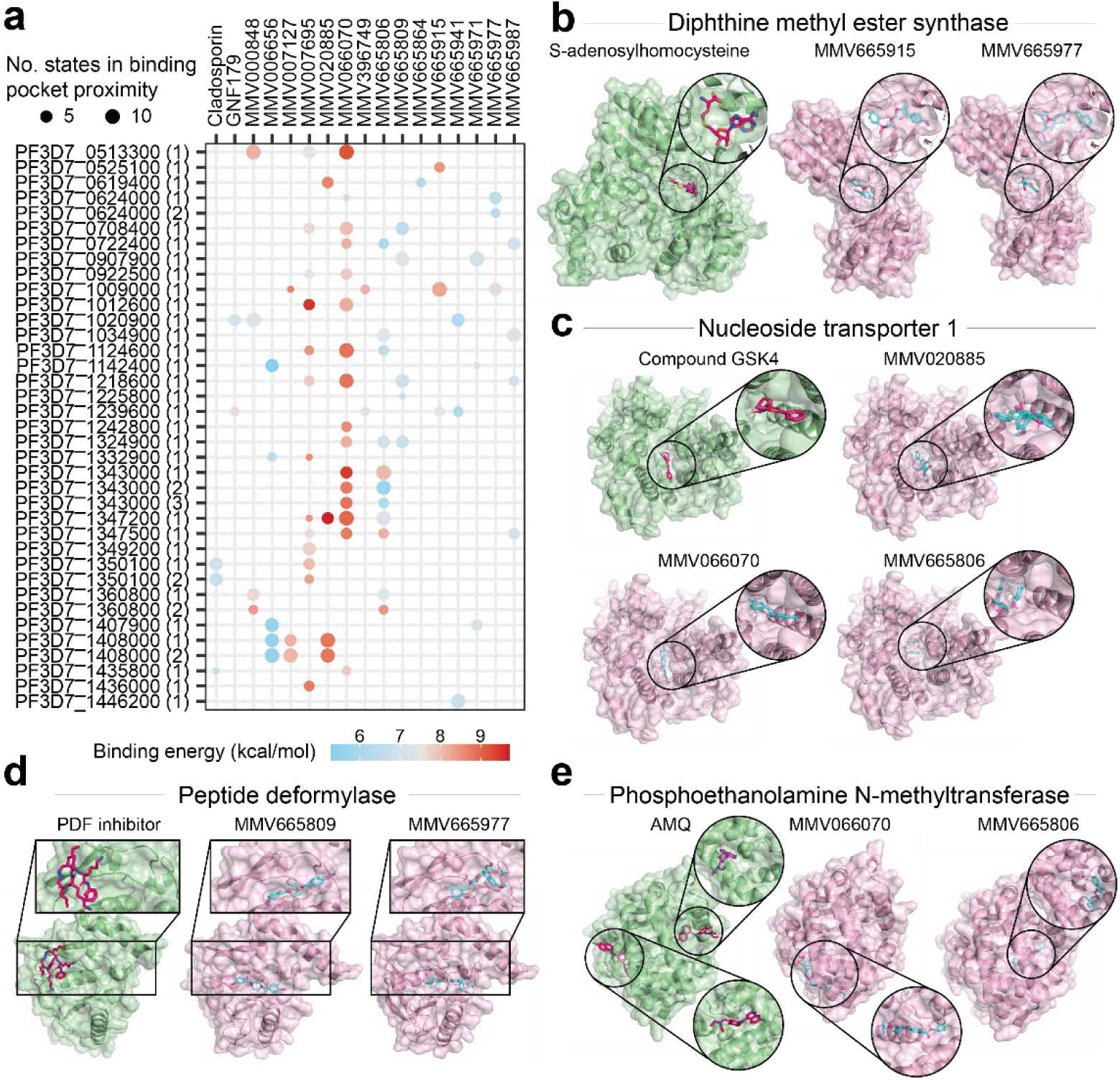
Docking screen of overlapping CETSA screen hits. **a** Docking results plotted on a grid chart. Mean estimated binding energy from docking is shown as point color from blue (low affinity) to red (high affinity) and number of conformations docked in the compared binding pocket (number in brackets after plasmoDB id) is shown as the point size. **b-e** Comparison of binding pocket of S-adenosylhomocysteine in DPH5 (**b**), GSK4 in ENT1 (**c**), PDF inhibitor in PDF (**d**) or amodiaquine in phosphoethanolamine N-methyltransferase (**d**) with compounds docked into the same binding pockets.

Besides these highlighted examples the in silico structure modelling suggested additional ligand-protein binding that may reflect antimalarial actions to previously studied protein targets including; (1) MMV666070 interacting with ethanolamine kinase (PF3D7_1124600)^47^; (2) MMV665941 binding to M17 leucyl aminopeptidase (PF3D7_1446200)^48^; (3) MMV665806 inhibiting lactate dehydrogenase (PfLDH, PF3D7_1324900)^49^; (4) MMV665977 binding to hexokinase (PF3D7_0624000)^50^; (5) MMV665806 and MMV665987 binding to methionyl tRNA synthetases^51^; (6) MMV666070 and MMV6658009 binding to arginyl tRNA synthetase^52^ (Supplementary Data 1). On the other hand, predicted ligand binding of glutamate tRNA ligase; Obg-like ATPase 1 (PF3D7_0722400) and coproporphyrinogen-III oxidase (PF3D7_1142400), have not been explored as potential antimalarial targets but warrant future studies (Supplementary Data 1). Obg-like ATPase is a promising target for antibiotic development for multiresistant bacterial species^53^.

### Detergent-assisted solubilization boosts membrane protein detection in P. falciparum

The current *P. falciparum* CETSA protocols, including the one used in the aforementioned screen, rely on standard protein lysate preparation methods compatible with quantitative proteomics measurements by mass spectrometry^23^. Due to the potential interference with the mass spectroscopy detections, these methods typically avoid the use of detergents for the extractions of membrane proteins. Indeed, most of the targets listed above are soluble proteins and the membrane proteins are underrepresented across all measurements (Extended Data Fig. 4a). For example, our initial screen failed to identify PfATP4, a known target of NITD246 and MMV1576856. To overcome this limitation for future applications, here, we wished to optimize the current protocol and incorporate detergent-based solubilization to expand the detection range to membrane-associated intracellular structures. This approach, termed detergent-assisted CETSA (^DA^CETSA), was first tested in both *lysate* and *in-cell* experiments, exposing either parasite lysates or cultures, respectively, to different concentrations of DHFR-TS inhibitor, pyrimethamine. For membrane solubilization, we opted for either 0.4% nonidet P-40 (NP-40) or 1% (n-dodecyl-β-D-maltoside and cholesteryl hemisuccinate) (DDM/CHS in 10:1 w/w) mixture. NP-40 has been previously used successfully in CETSA of mammalian cells ^54–56^ and the DDM/CHS mixture is frequently used for the extraction of folded membrane proteins for structural biology applications^57–59^. Subsequently we incorporated the SP4 protein extraction protocol that mediates efficient removal of all contaminants including detergents ^60^ (Fig. 4a and material and methods).

**Figure 4.**
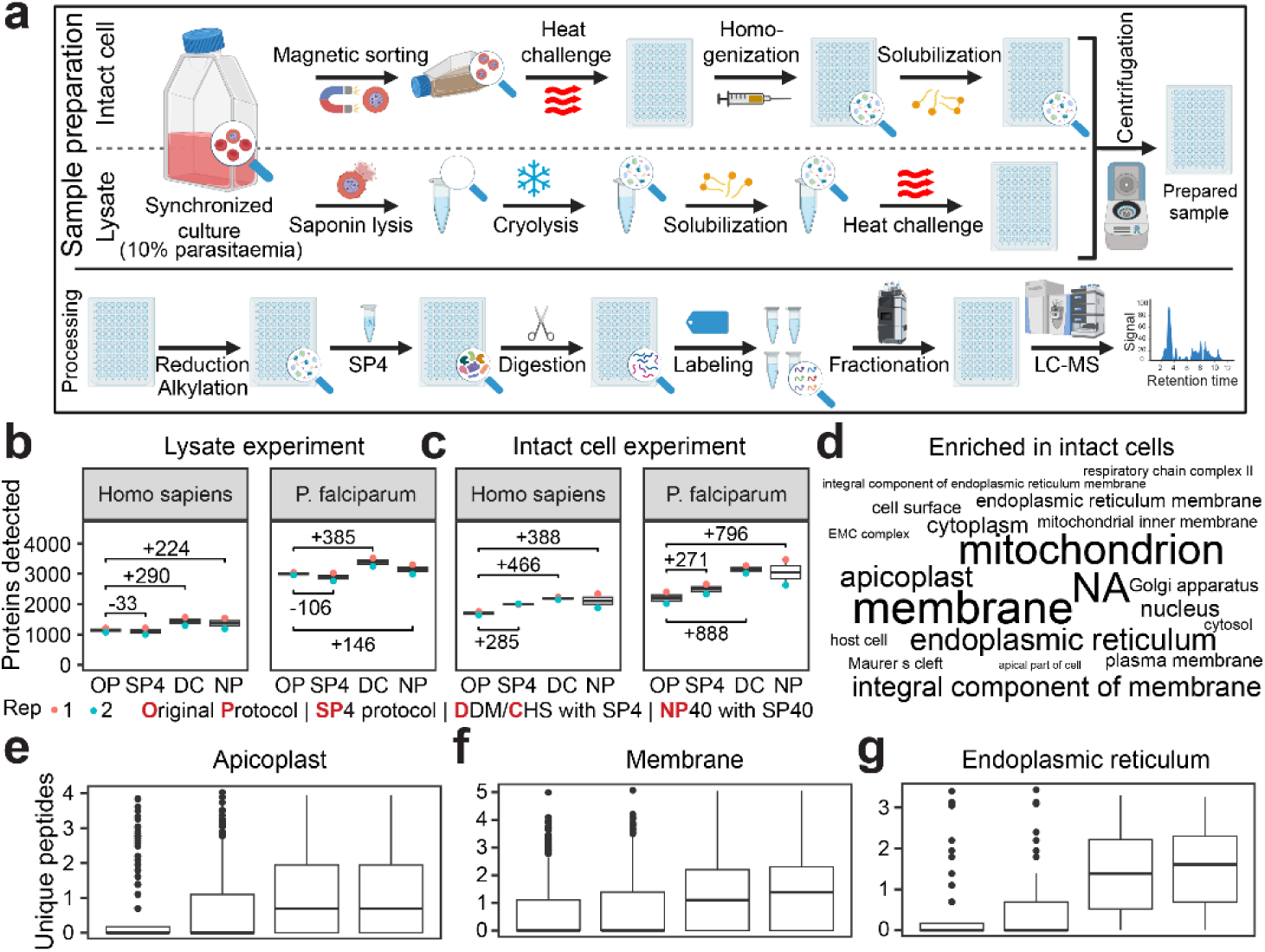
Detergent solubilization boosts up detection of membrane proteins. **a** ^DA^CETSA pipeline includes a solubilization step after cryolysis (lysate setup) or cell homogenization (intact cell setup). SP4 cleanup follows reduction and alkylation prior to the digestion. The additional steps are marked in bold. **b-c** Comparisons of the numbers of detected human (Homo sapiens) and parasite (P. falciparum) proteins from cell lysates (**b**) and intact cells (**c**) using different protein extraction protocols: original protocol (OP), SP4 without detergent (SP4) and SP4 with detergents DDM/CHS (DC) or NP40 (NP). The experiments were performed in duplicates and the numbers above the boxplots indicate the mean increase in the number of detected proteins. **d** Word cloud representations of cellular component GO terms of proteins that were uniquely detected only in the detergent conditions from intact cell samples. The font size correlates with the number of proteins with the associated GO component term. **e-g** Comparisons of the numbers of unique peptides detected in samples using different extraction protocols across duplicate experiments. The proteins are grouped by GO component terms, showcasing apicoplast proteins (**e**), membrane-associated proteins (**f**) and proteins localised to endoplasmic reticulum (**g**). Panel A was created with BioRender.com

Compared to the standard CETSA protocol, DDM/CHS or NP40 solubilization yielded on average additional 358 and 146 parasite proteins in *lysate* ^DA^CETSA and 888 and 796 parasite proteins in *in-cell* ^DA^CETSA experiments, respectively (Fig. 4b-c). There were also significant increases in the numbers of human red blood cell proteins detected by both types of experiments (+290 and +224 in *lysate*, +466 and +388 in *in-cell* experiments for DDM/CHS and NP40, respectively). Functional analyses using gene ontology (GO) revealed that ^DA^CETSA in both *lysate* and *in-cell* experiments expanded the detection of proteins localized to membrane-rich organelles such as endoplasmic reticulum, apicoplast, and mitochondria, but also of exported structures in the host cell cytoplasm including Mauer’s clefts (Fig. 4d, Extended Data Fig. 4d). These also included proteins without an assigned GO component (non-annotated proteins, NA) indicating that a considerable proportion of functionally uncharacterized, unique *P. falciparum* proteins are associated with membranes or localized in membrane-rich organelles. For the most overrepresented functional protein groups, we also observe an increased number of identified unique peptides derived from the samples solubilized by both DDM/CHS and NP40 (Fig. 4e-g). These results suggest that detergent-assisted solubilization massively improves the protein coverage of mass spectrometry detection in *lysate* and *in-cell* CETSA by expanding detection of membrane-associated proteins.

### ^DA^CETSA identifies soluble and membrane protein drug targets

First, we verified that the use of detergents in ^DA^CETSA does not prevent the identification of soluble targets in *P. falciparum*. Indeed, for pyrimethamine, PfDHFR-TS was the most stabilized protein in *lysate* experiments using both DDM/CHS and NP40 (Fig. 5a-c). In the *in-cell* experiments, DHFR-TS also displayed thermal stabilization upon pyrimethamine exposure, albeit with somewhat lower confidence (Extended Data Fig. 4e). Similarly, *lysate* ^DA^CETSA with two experimental antimalarials, DSM705 and DSM265^61,62^, uncovered *P. falciparum* dihydroorotase dehydrogenase (PfDHODH, PF3D7_0603300), the expected protein target of both compounds (Fig. 5d-e). These results indicate that the presence of detergents did not impact genuine drug interactions with their soluble targets. Curiously, *lysate* ^DA^CETSA for pyrimethamine as well as DSM705 and DSM265 detected confident thermal shift for additional protein sets (outside their presumed targets) that did not overlap with those detected by the standard protocol (Fig. 5a-e, Table 4). Currently, it is unclear if these proteins represent additional pyrimethamine targets or an off-target binding. Next, we applied ^DA^CETSA to detect membrane-associated protein targets, by investigating MMV1576856, a racemic mixture of cipargamin known to target endoplasmic reticulum membrane-associated PfATP4 ^63^. Indeed, PfATP4 was the top target candidate of MMV1576856, in both *lysate* and *in-cell* experiments (Fig. 5f-g). In the *in-cell* experiment, MMV1576856 induced stabilization of a large number of proteins, some of which may represent additional targets, while others reflect a strong cellular response to the treatment via downstream effects. GSEA of these thermal shifts identified protein factors related to transcription and translation (Fig. 5h). Interestingly, while the ribosome and T-complex were significantly destabilized, proteins related to translation initiation and its regulation were stabilized, indicative of ribosome stalling. These results hence indicate that while cipargamin indeed binds PfATP4 and its cytotoxic effect on the parasite cell is exerted via disruptions of gene expression and protein turnover. in the final stage of this study, we carried out ^DA^CETSA with atovaquone that is known to binds to the Rieske protein, a subunit of membrane mitochondrial respiratory complex III (bc1 complex) ^64^. Indeed, in the *lysate* experiment, we identified 2 out of 9 detected bc1 complex subunits as the top hits. Although Rieske protein displayed concentration-dependent stabilization as expected, the shift did not pass the confidence cutoff, reflecting the intrinsic protein properties in our experimental setup rather than the lack of binding (Fig. 5i). In the *in-cell* experiment, all 7 detected bc1 complex subunits were significantly stabilized among a multitude of other proteins (Fig. 5j). GSEA showed that atovaquone treatment stabilized not only respiratory complex III (the direct target) but proteins integral to membrane in general. On the other hand, citrate cycle and the synthesis of secondary metabolites were among the destabilized terms (Fig. 5k). Surprisingly, the strongest hit in the lysate atovaquone data was not the respiratory complex III but a putative monocarboxylate transporter PF3D7_0926400 (Fig. 5l), recently labelled as MCP2^65^. MCP2 was also stabilized in the intact cells, albeit with lower confidence (Extended Data Fig. 5e). To verify its stabilization, we performed proteome integral solubility alteration (PISA), an assay built on the same principles as CETSA^66^. Performed with parasite lysates, PISA identified MCP2 as the only hit (Fig. 5m, Supplementary Table 6) with almost doubled solubility compared to the control or DSM265-treated samples (Fig. 5N, Extended Data Fig. 5f). These results suggest that MCP2, aside from its native function, can bind and possibly transport atovaquone.

**Figure 5.**
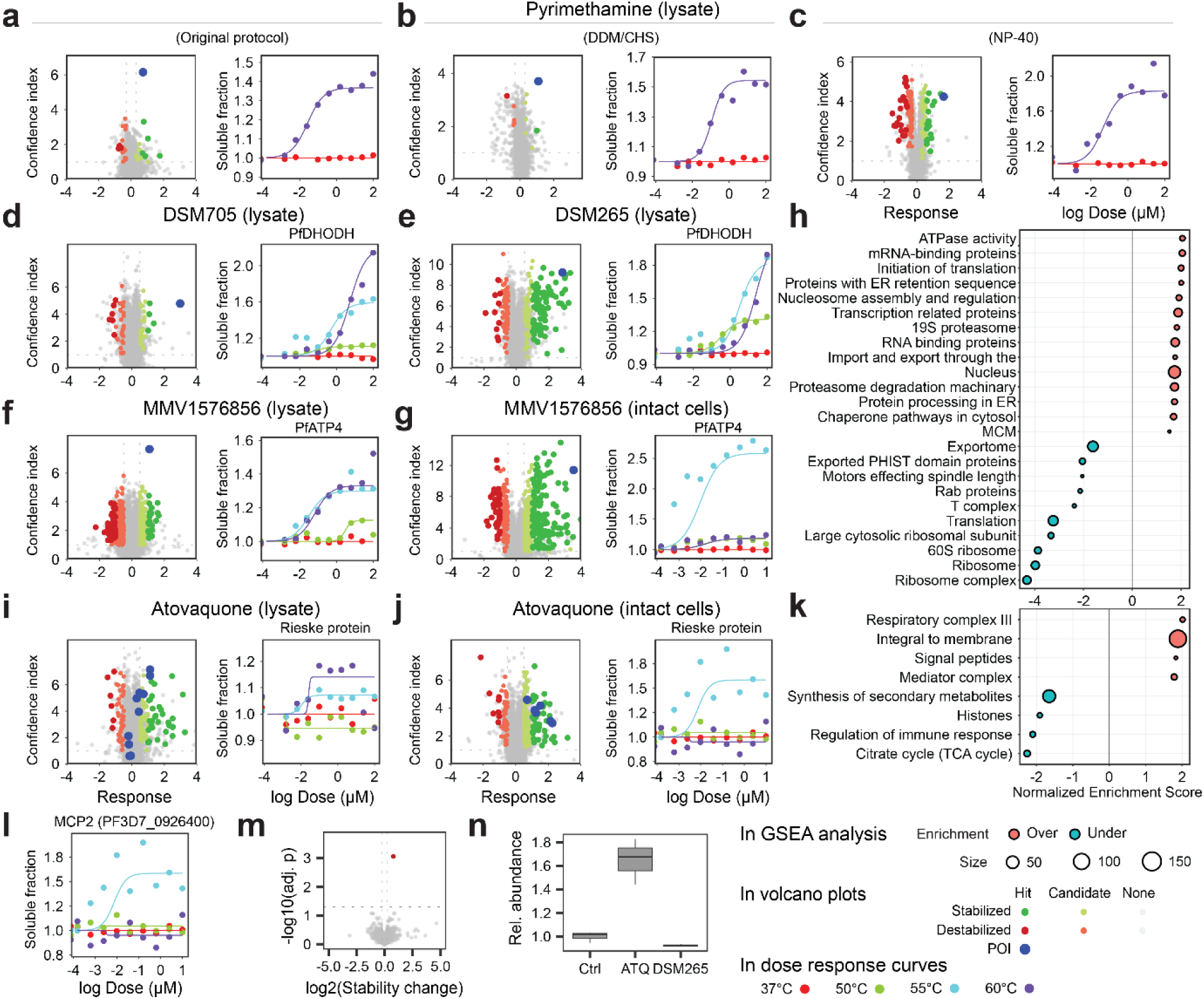
Detergent solubilization is compatible with target identification in P. falciparum. **a-c** Left: volcano plots show correct identification of DHFR (blue) as the target of pyrimethamine with the original CETSA protocol (**a**) as well as with the improved protocol that includes detergent solubilization (**b** DDM/CHS or **c** NP40) and SP4 protein extraction/ Right: fitted melting curves of DHFR with increasing doses of pyrimethamine. **d-i** Volcano plots on the left and dose-response curves on the right shown for the selected drugs and their targets: DSM705 and PfDHODH (**d**), DSM265 and PfDHODH (**e**); atovaquone and Rieske protein in the lysate (**f**) and intact cell (**g**) setup; MMV1576856 and PfATP4 in the lysate (**h**) and intact cell (**i**) setup. **j-k** Gene set enrichment analysis using intact cell data from atovaquone (**j**) and MMV1576856 (**k**) challenge. **l** Isothermal dose response curves of MCP2 upon atovaquone treatment. **m** Volcano plot of PISA experiment, showing MCP2 as the strongest stabilized protein. **n** Relative abundance of MCP2 was ∼1.7x higher compared to the mock control and DSM265-treated sample.

## Discussion

Since the conception of CETSA in 2014, proteome-wide stability assays have become instrumental in deconvolving drug MoA. Here, we screened 25 drugs with known or unknown MoA by *lysate* CETSA, with the goal of identifying their drug targets. Identifying several known drug targets like lysyl-tRNA ligase (cladosporine), PfPNP (MMV000848) and falcilysin (MMV00848 and MMV665806) and validating ACS10 as the target of MMV665915 gives us confidence in the results of the screen. Interestingly, some compounds stabilized well-studied proteins and drug targets (Fig. 1), including plasmepsins I and II. Although plasmepsins I and II are thought to be disruptable off-targets of compounds that target essential plasmepsins (reviewed in ^67^), we did not detect stabilization of other plasmepsins in these datasets. It is likely that in the lysate setup, the compounds stabilizing plasmepsins are more likely to bind to more abundant off-targets, masking the engagement with the real targets (potentially plasmepsins IX or X). *In vivo*, these drugs might not localize in the digestive vacuole and not bind these off-targets at all. Another well-known drug target that was stabilized by two compounds was PfHsp90. PfHsp90 inhibitors have been previously identified from a screen of pharmaceuticals and FDA-approved drugs, and they were found to inhibit the parasite growth^41–43^.

The proteome-wide assays often reveal a more complex picture of cellular drug interactions than previously thought. For example, the median number of hits in a large screen performed with compounds targeting human cell targets by *Vranken et al* was 9.5^68^, which is comparable with our median of 3 hits in 4x smaller *P. falciparum* proteome. The authors showed that some of the identified proteins are off-targets. Off-targets are certainly also present as hits in our datasets. Although the main drug targets were often the top hits (Fig. 1b,H, Fig. 4a,d-e, Fig. 5f, Extended Data Fig. 2a-b), their ranking depends on the ability of the individual proteins to be stabilized by drug binding^25^. As a result, the drug targets can be ranked lower than off-targets or not be significantly stabilized at all (false negatives). Indeed, the hit rate of compounds with known MoA in the study by *Vranken et al* was 73%.

Another potential culprit in CETSA screens are false positives. We took two measures to reduce their numbers in our screen. First, we developed *ITDRMS* pipeline which removes some of the false positives that are not excluded by our previous package, *mineCETSA*. Second, we further filtered the hits using molecular docking, in our case effectively narrowing down the main hits to 13 proteins (Fig. 3). Of these, DPH5 was the top hit for here compounds and could also be disrupted in PiggyBac screen ^69^, highlighting their potential as drug targets. ENT1 is, interestingly, plays a role in the same pathway as PfPNP as it transports the PfPNP product, hypoxanthine, from parasitophorous vacuole into the cytosol^70^. ENT1 has recently been shown to be druggable and its cryoEM structure with a bound inhibitor has been solved^71^. PMT is an interesting drug target as its inhibitors with limited IC50 values have previously been found^46,72^. PDF is a well-studied drug target in bacteria as human cells lack its homolog (reviewed in ^73^), but the inhibitors originally associated with PDF were found to target a different protein in *Plasmodium*. Not much available literature on *Plasmodium* ethanolamine kinase. While it was suggested to be the target of artemisinin^47^, it remains uncharacterized both as a protein and as a drug target. Of note, we could not evaluate 17 proteins that lack homologous structures and could represent equally plausible drug targets (Supplementary Table 3).

Next to off-targets and possible false positives, our screen showed stabilization of three membrane proteins (Extended Data Fig. 4a-c) which are not necessarily drug targets but could play a role in their transport across membranes. Indeed, mutations in transporters PfCRT and PfMDR1 confer resistance to a number of clinically used drugs. This has motivated us to add the membrane solubilization step to our sample preparation, leading to subsequent validation of mitochondrial complex III and PfATP4 as the targets of atovaquone and cipargamin, respectively, without impacting the identification of soluble drug targets (Figures 4 and 5). Surprisingly, one of the strongest hit in CETSA assays and the only significantly stabilized protein in lysate PISA assay by atovaquone was the monocarboxylate transporter, MCP2 (Fig. 5L-N). MCP2 has been originally predicted to localize in the apicoplast, however, the recent hyperLOPIT screen could not predict a clear localization^74^. We hypothesize that MCP2 at least partially localizes to the mitochondrial membrane where it transports atovaquone to the site of action. Further experiments will be necessary to confirm this finding.

Finally, although we initially used CETSA with lysate samples to interrogate direct drug-protein interactions, we also tested ^DA^CETSA on intact cell samples to validate the membrane protein targeting drugs. In intact cells, atovaquone and cipargamin stabilized not only their respective targets, but also a number of other proteins, reflecting the downstream metabolic effects and stress responses (Fig. 5H,K). Interestingly, the response to atovaquone seemed more specifically connected to its main target, respiratory complex III. The downstream metabolic effect was the destabilization of the TCA cycle, which we interpret as the stalling of TCA cycle caused by reduced recovery of redox cofactors due to the inhibition of the electron transport chain. The products of TCA cycle are used for the synthesis of secondary metabolites, which was another destabilized term. Additionally, the stabilization of the mediator complex and the destabilization of histones suggest an initiation of transcriptomic response of the parasites after 1h of treatment. Contrary to atovaquone, cipargamin shows a larger and less specific response, in agreement with its fast-acting nature. Interestingly, large ribosomal subunit was significantly destabilized while the proteins connected to initiation of translation were stabilized. This indicates inhibition of the translation initiation and subsequent formation of stress granules (reviewed by *Buchan and Parker*^75^). The stress granules are heterogeneous particles that, among other proteins, typically contain translation initiation factors, RNA-binding proteins, chaperons and MCM complex subunits, all of which are stabilized by cipargamin in intact cells. Stress granules also contain 40S ribosomal subunits but not 60S ribosomal subunits, which were destabilized by cipargamin. On top of that, cipargamin induced stabilization of 19S proteasome and proteasome degradation machinery, which together with formation of the stress granules indicates a complete dysregulation of protein homeostasis.

Altogether, we identified a number of candidate drug targets of screened compounds, combining an experimental (CETSA) and computational (docking) approach. Building on our original protocol, we took steps to improve the identification of membrane protein targets both in lysates and intact cells and validated the MoA of atovaquone and cipargamin, additionally mapping their effect on live parasites.

## Acknowledgements

We would like to thank Jeremy Burrows and Didier Leroy (Medicines for Malaria Venture, Geneva, Switzerland) and Amit Sharma (Cladosporin) for providing us with some of the compounds used in this study. This work was supported by Singapore Ministry of Education grant #MOE2019-T3-1-007 and Singapore National Science Foundation grant #NRF-CRP24-2020-0005 (ZB); (ZB) and NTU-PPF-2019 (JMD).

## Data availability

The raw mass spectrometry files were deposited to PRIDE under accession numbers: PXD048737, PXD048745, PXD048750, PXD048752, PXD048756, PXD048757, PXD048761, PXD048768 and PXD048772.

## Extended Data Fig.s

**Extended Data Fig. 1.**
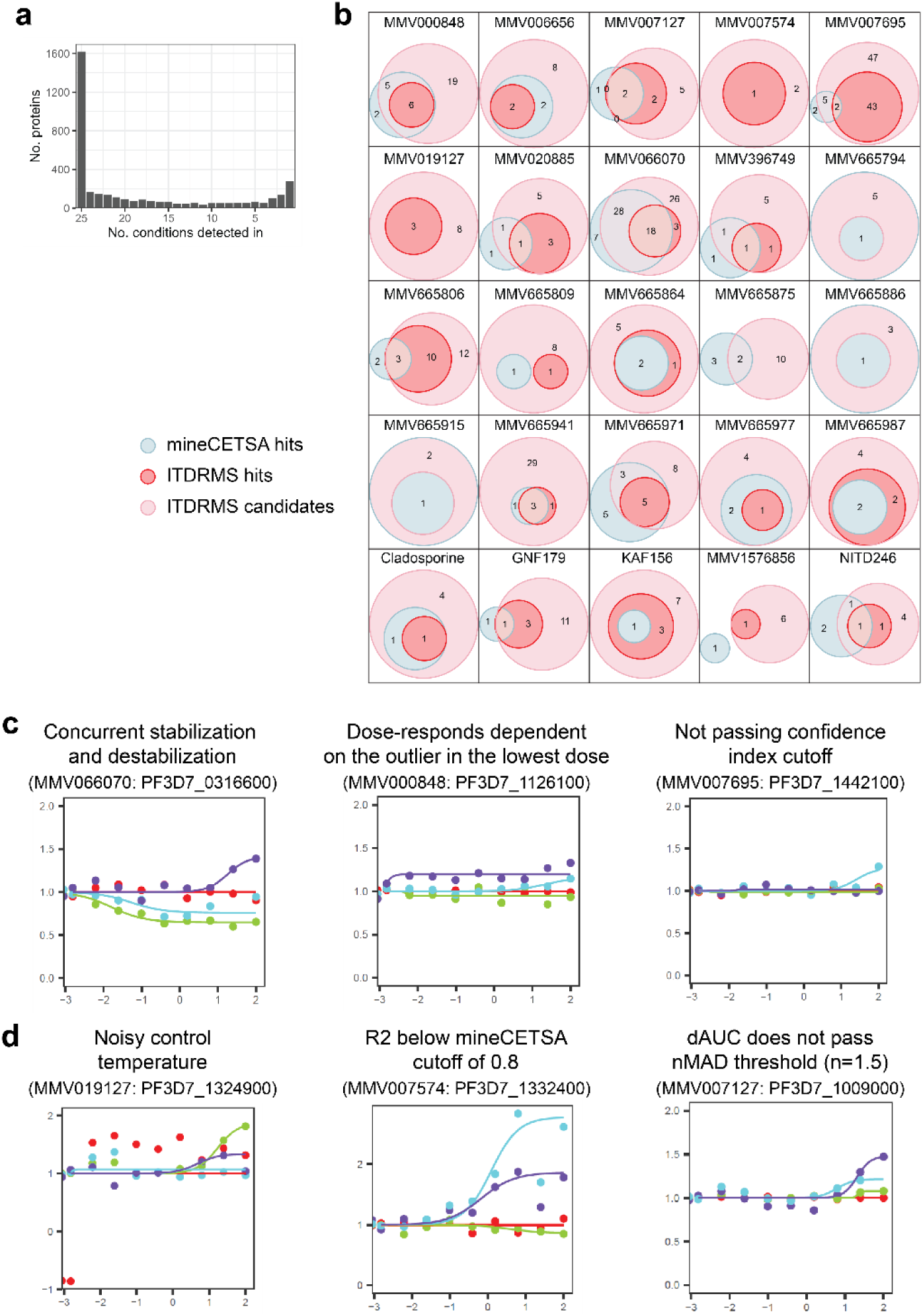
Comparison of *mineCETSA* and *ITDRMS* packages for data analysis. **a** Number of proteins detected in all 25 or less experiments in the drug screen. **b** Ven diagrams showing overlaps in hits identified by the two packages. **c** Examples of curves that were identified by mineCETSA but not *ITDRMS*. **d** Examples of curves that were identified by *ITDRMS* but not *mineCETSA*.

**Extended Data Fig. 2.**
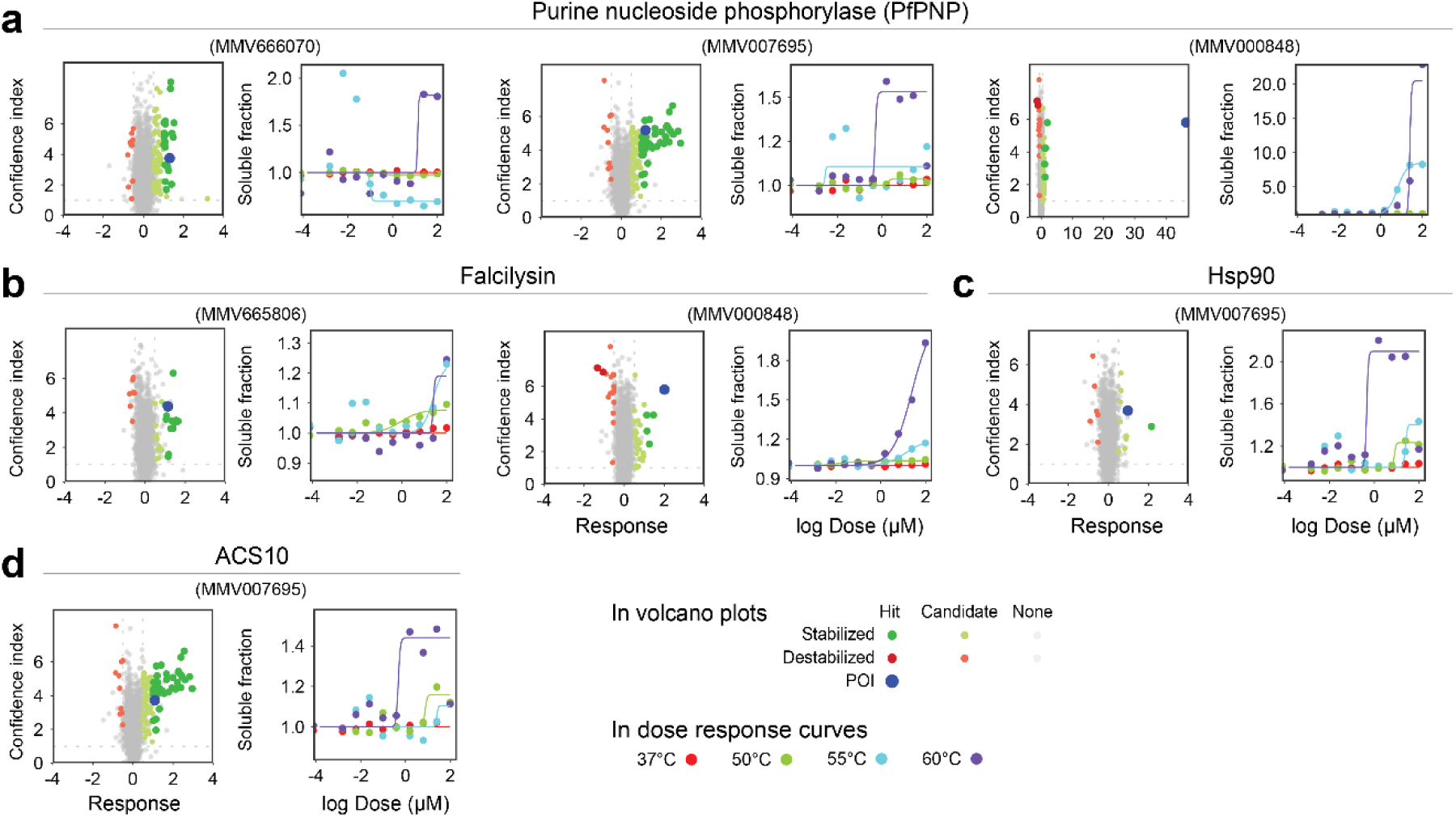
Result of the CETSA screen. **a-d** Volcano plots (left) and dose-response curves (right) of hit-compound pairs**. a** MMV compounds stabilizing purine nucleotide phosphorylase. **b** MMV compounds stabilizing falcilysin. **c** MMV007695 stabilizing Hsp90. **d** MMV007695 stabilizing ACS10. **e** Volcano plots of three MMV compounds that stabilize a large number of ribosomal subunits and translation factors. **f** Volcano plots of three MMV compounds that stabilize subunits of T-complex.

**Supplementary Figure 3.**
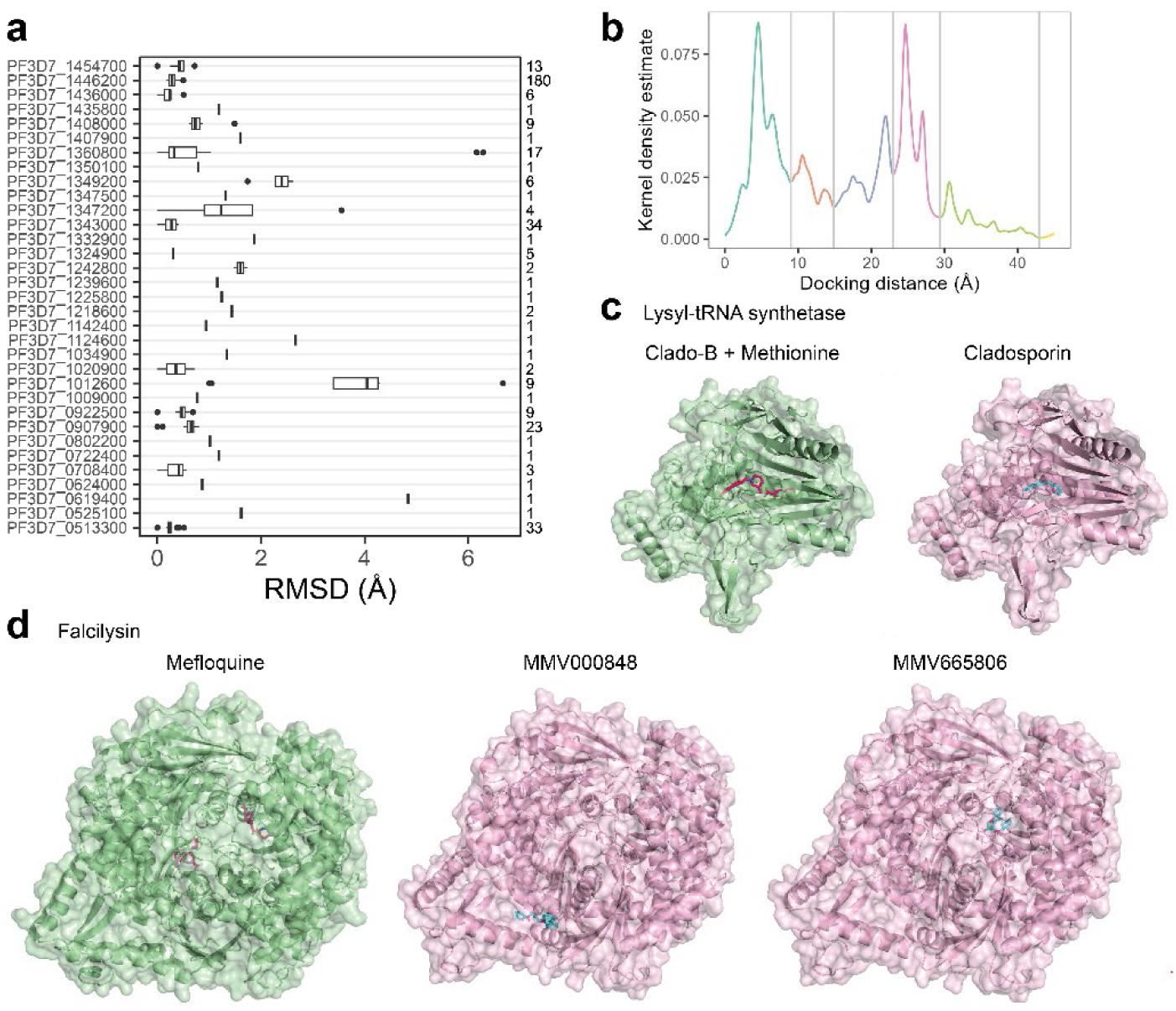
Docking results. **a** Root mean square deviation (RMSD) of structures of control proteins with bound ligands to the docked proteins. The numbers on the right denote the number of control protein structures. **b** Distribution of the distances of docked compounds from the ligands in control proteins, clustered into five groups that are denoted by different colors. **c-d** Comparative visualization of structure of control proteins with bound ligands (green) and proteins docked with the tested compounds. **c** Lysl-tRNA synthetase. **d** Falcilysin.

**Extended Data Fig. 4.**
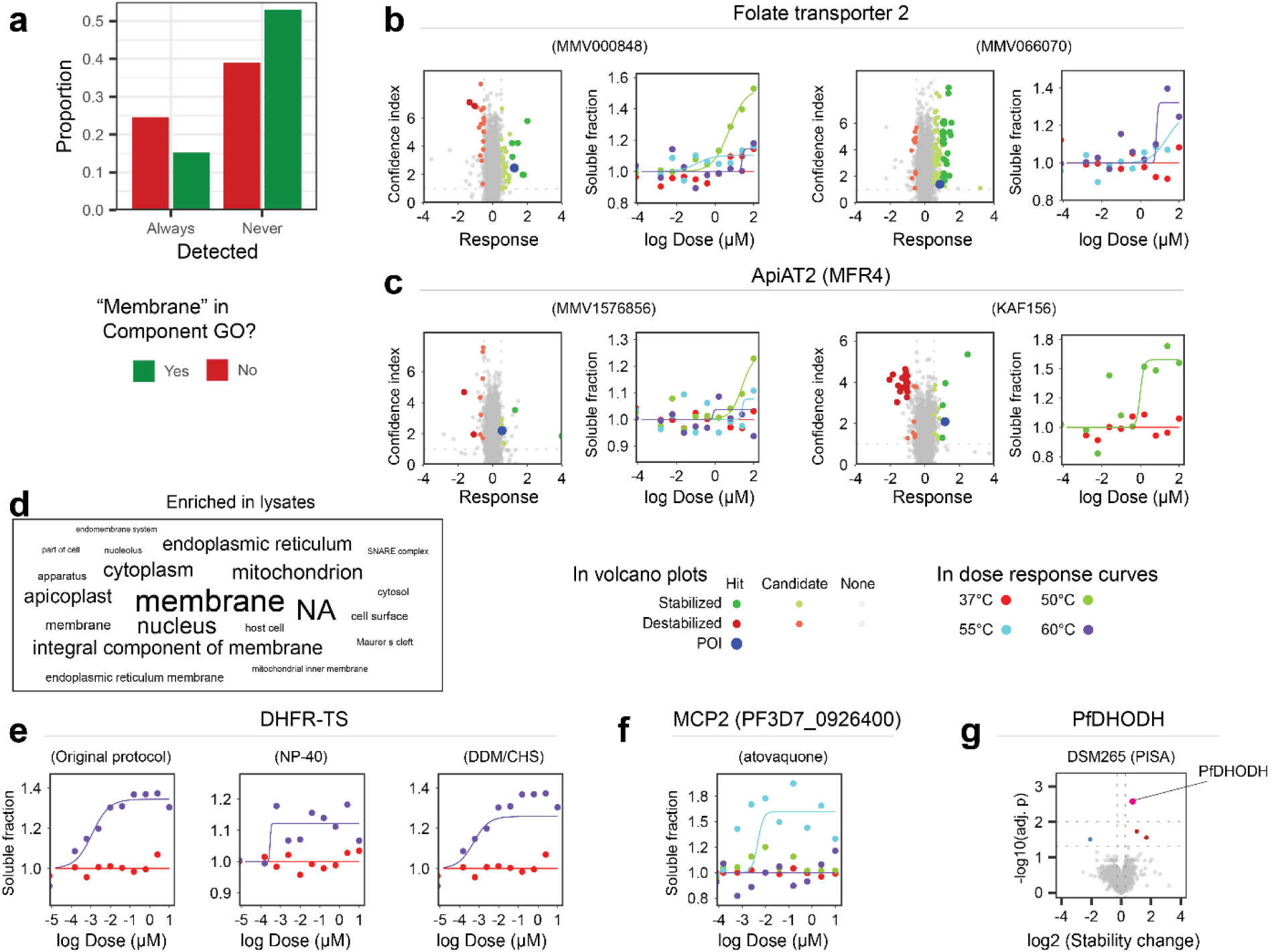
daCETSA results. **a** Only 16% of all membrane proteins was detected in all measurements of the screen, compared to 25% of all non-membrane proteins. Conversely, 53% of all membrane proteins were never detected, compared to 39% of non-membrane proteins. **b-c** Volcano plots (left) and dose-response curves (right) of membrane proteins stabilized in the initial screen. **b** Folate transporter 2 stabilized by DHFR inhibitors MMV000848 and MMV666070. **c** ApiAT2 (MFR4) stabilized by MMV1576856 and KAF156. **d** Wordcloud representation of the terms enriched in the lysate samples prepare with the assistance of detergents. **e** Dose-response curves of DHFR-TS stabilized by pyrimethamine using the original protocol, SP4 with NP-40 and SP4 with DDM/CHS in the intact cells. **f** Stabilization of MCP2 in the intact cells by atovaquone. **g** PfDHODH identified as the hit by PISA with DSM265.

## Experimental Procedures

### Experimental Design and Statistical Rationale

The screen CETSA experiment was conducted exactly as previously described in our protocol for CETSA optimized for *P. falciparum*^23^. The *ITDRMS* analysis pipeline has been compared to existing mineCETSA pipeline^23^ and described in this text. The R package *ITDRMS* is available on GitHub^76^. Proteins with confidence index < 0.01 and response >1 were considered hits. Proteins with -log10(confidence index) < 0.05 and response >0.5 were considered candidates. The results of PISA assay were analysed by empirical Bayesian model. Proteins with adjusted p-value <0.05 and log2FC > 1.2 were considered hits. Molecular docking is described in this text and the approach to its analysis is explained in the results.

### Parasite cultures

*P. falciparum* 3D7 cultures were maintained in RPM-1640 medium (Gibco) supplemented with 0.25% Albumax II (Gibco), 0.1 mM hypoxanthine (Sigma), 0.2% sodium bicarbonate (Sigma), and 10 mg/L gentamycin at 37°C with 5% CO_2_, 3% O_2_, and 92% N_2_ gas mixture and kept agitating at 37°C. Media was changed daily while the haematocrit was set at 2% and parasitaemia kept below 10%.

### Intact cell sample preparation

Haematocrit of well synchronized parasite culture at trophozoite stage (32-38 hpi) was adjusted to 10%. The culture was loaded on MACS CS column (Miltenyi Biotec), washed with culture media and the enriched trophozoites (>80% parasitaemia) were eluted. The trophozoites were recovered at 2% haematocrit at 37°C incubation with agitation for 1 hour before drug challenge. The culture was split into 10 wells (10 million cells per well) and treated with 4-fold dilution series of the drug (starting at 10mM and with DMSO as control. After 1 hour, the cells were pelleted, washed with 1x PBS and resuspended each in 100 μL PBS. The samples were split by 25 μL and each set of split samples was subjected to thermal challenge for 3 min (37°C and 65°C for pyrimethamine experiments, 37, 50, 55 and 60°C or 37, 51 and 57°C in the initial drug screen, and 37, 53, 59 and 65°C when detergents were used). The samples were cooled to 4°C immediately following the thermal challenge, diluted in 1:1 ratio with lysis buffer (50 mM HEPES pH 7.5, 5 mM beta-glycerophosphate, 0.1 mM 25 Na_3_VO_4_, 10 mM MgCl_2_, 2 mM TCEP, and cocktail EDTA-free protease inhibitors (Sigma) and flash frozen. The cells were subsequently subjected to three freeze-thaw cycles, sheared by 10 passes through 26-gauge needle and centrifuged (20,000 x g, 20 min, 4°C) and the supernatant was recovered and flash-frozen.

### Lysate sample preparation

Harvested parasite pellets were cryolysed (Precellys® Evolution CryoLyser) in 6 cycles with 5s shaking at 6000 RPM and 25s break at 4°C. For detergent solubilization, 1%/0.1% (w/v) DDM/CHS or 0.4% NP40 were added to the lysed pellet and incubated at 4°C rotating for 1 hour. The samples were centrifuged (20,000 x g, 20 min, 4°C) and the supernatants were collected. The protein concentration was measured with BCA™ protein assay kit (Thermo Scientific). For CETSA, 100 μL of 2.5 mg/ml lysate each were added to 9 tubes of 4-fold drug dilution series starting at final concentration of 100 μM and to 1 tube with DMSO control. Each tube was split to 4 tubes of 10 samples with 25 μL volume each and after 3 minutes of incubation (room temperature), these were subjected to heat challenge at 37, 50, 55 and 60°C for 3 min and cooled down to 4°C. For PISA, the lysates were treated with 10μM compound or DMSO control in triplicates for 5 minutes, then each sample was split into 12 parts that were subjected to heat challenge at 50-72°C with 2°C increment and pooled together. The samples were centrifuged (20,000 x g, 20 min, 4°C) and supernatants were collected for downstream processing.

### ^DA^CETSA sample preparation

^DA^CETSA samples were prepared as described above with the following modification: 1%/0.1% (w/v) DDM/CHS or 0.4% NP40 was added to the sample and incubated for 1h at 4°C on rotatory wheel after the mechanical lysis before the centrifugation at 20,000 x g. Both detergents were tested separately in experiments with pyrimethamine. Subsequently, DDM/CHS was used for lysate CETSA and NP40 was used for intact cell CETSA experiments.

### Sample processing

20 μg of protein was incubated in a denaturation and reduction buffer (TEAB, pH 8.5), 20 mM tris(2-14 carboxyethyl)phosphine) (TCEP, pH 7.0), and 1% (w/v) Rapigest (Waters)) at 55°C for 20 minutes. 55 mM of CAA was added and incubated at room temperature in the dark for 30 minutes. For the initial screen, TFA was added to 1%, samples were incubated (45 min, 37°C, shaker) and centrifuged (20,000 x g, 15 min). The supernatants were dried in centrifugal vacuum evaporator, washed with 200mM TEAB (pH 8.5) and resolubilized in 100mM TEAB to 1 μg/μL concentration. For SP4-based protocol,1 mg of glass beads and 150 μL of ACN were added to each sample. Samples were centrifuged (16,000 x g, 5 min), and the supernatant was removed carefully without disturbing the beads. Samples bound to the beads were washed 3x with 250 μL of 80% ethanol, resuspended in 70 μL of 100 mM TEAB with Lys-C (4.6 AU, Wako) and trypsin (1 μg), and incubated for 18 hours at 37°C with agitation. After a short spin to pellet the beads, the supernatants were collected and dried by a centrifugal vacuum evaporator. Samples were resolubilized in 100mM TEAB to 1 μg/μL concentration.

### TMT labelling and fractionation

8 μg of digested peptides were labelled with 4ul TMT10plex Isobaric Label Reagent Set (Thermo Fisher Scientific) incubated in room temperature overnight before it was quenched with 1M Tris (pH 7.4). The labelled samples were pooled and desalted using Oasis HLB column (Waters), eluted solution was dried. The samples were resolubilized in 5% (v/v) acetonitrile, 5% (v/v) ammonia and run through high pH reverse phase liquid chromatography column (Zorbax 300 extend C-18 4.6mm x 250mm). Collected fractions were concatenated into 20 fractions and dried.

### Mass spectrometry and protein quantification (CETSA)

The dried peptide samples were reconstituted in a solution of 1% acetonitrile, 0.5% (v/v) acetic acid, and 0.06% trifluoroacetic acid (TFA) in water. For CETSA, the samples were analyzed on Dionex 3000 UHPLC system coupled with a Q Exactive HF mass spectrometer (Thermo Fisher Scientific). Peptides were separated on 50cmx75μm analytical column (EASY-Spray™, Thermo Scientific) over a 70-minute gradient (1-55 min: 2-25%, 55-57 min: 25-50%, 57-58 min: 50-85%, 58-63 min: 85%, 63-70 min: 2%). Mass spectrometric data were acquired using data-dependent acquisition (DDA) with full scan MS spectra (350-1550 m/z, resolution: 70,000, AGC target: 3e6) and Top12 MS2 spectra (resolution: 35,000, AGC target: 1e5, 1e5 isolation window at 1.2 m/z). Proteome Discoverer 2.1 software (Thermo Fisher Scientific) was used for protein identification and quantification based on Xcalibur raw files. Sequest HT (Thermo Scientific) search engine used P. falciparum 3D7 sequences from plasmoDB (version 66) and Uniprot database for human proteins. Parameters were as follows: 30 ppm MS precursor mass tolerance, 0.06 Da fragment mass tolerance, a maximum of 3 missed cleavage sites. Dynamic modifications used were oxidation (M), deamidation (NQ), and acetylation (N-terminal protein) and the static modification was carbamidomethylation (C). False discovery rate (FDR) estimation employed forward/decoy searches, with acceptance criteria set at high FDR (1%) and medium FDR (5%) levels for peptide and peptide spectrum matches (PSMs).

For PISA, samples were analyzed using EASY-nLC 1200 coupled to an Orbitrap Exploris 480 mass spectrometer (Thermo Fisher Scientific). Peptides were separated on a reversed-phase column using a linear gradient of 3–28% ACN in 0.1% FA over 90 min, 28-40% ACN over 5 min and 40-80% ACN over 5 min at a flow rate of 200 nL/min. Full scan range of m/z 350–950 at a resolution of 45,000 was followed by 34 DIA scans in custom-defined windows covering m/z 145–1450 with an Orbitrap resolution of 22,500. The normalized AGC target was set to 300 and 3000 and maximum injection time was set to 99.61 and 40.6ms for MS1 and MS2, respectively. HCD fragmentation was applied with normalized collision energy of 27%. The RAW files were converted to mzML using msconvert^77^ and the spectra were analyzed using DIA-NN^78^ with 2 allowed missed cleavages, maximum number of variable modifications set to 2, allowed N-terminal methionine excision, cysteine carbamidomethylation, methionine oxidation, N-terminal acetylation, peptide length range set to 6-30 and using our in-house *P. falciparum* spectral library.

### CETSA analysis

CETSA analysis was performed using in-house-developed package *ITDRMS* available on GitHub^76^. The protein abundances were scaled based on the sum of the abundances from *P. falciparum* proteins only. The mean of first two data points (lowest concentration and DMSO control) at each temperature were normalized to 1. The data were then normalized by median ratio value across each condition. For the baseline data at 37°C, one outlier point was removed when the fit *y=a+x* to the data with that point missing had significantly increased R^2^ compared to fits to data missing no points or missing any one of other points. R^2^ was considered significantly increased when the z-scaled R^2^ value was larger than or equal to 2. The baselines were re-normalized to 1. The baseline data were then fitted with *y=1+a* and thermal challenge data were fitted with log-logistic function with R package minpack.lm^79^ with one allowed removal of outlier point as described. The confidence intervals of the linear (baseline) and log-logistic (thermal challenges) fits are then used to calculated the confidence index for each thermal challenge temperature and each protein as 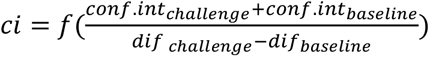, where *dif* is difference between the ratio value at thermal challenge and baseline temperature, *conf.int* are the respective confidence intervals and function *f()* uses log-logistic fit to transform the values to range from 0-1, resembling p-values. *ci* values are adjusted Benjamini-Hochberg method across each challenge temperature and multiplied across each protein. The total response is calculated as the maximum response (out of all thermal challenge temperatures) plus the sum of responses at each thermal challenge temperature. Response is defined as the highest value of the fit for stabilizing curve or the lowest value of the fit for destabilizing curves.

### PISA

Triplicates of detergent-solubilized *P. falciparum* lysates treated with atovaquone, DSM265 or DMSO were split into PCR tubes and heated to 12 different temperatures (51-73°C, 2°C interval) for 3 minutes, cooled down to 4°C and equal volumes were pooled back into one tube per replicates, centrifuged (20,000 x g, 20 min, 4°C). Equal volumes of supernatants were processed using SP4 protocol described above and the peptides acidified with 0.1% FA. 1 μg of peptides of each sample were injected using EASY-nLC 1200 coupled to an Orbitrap Exploris 480 mass spectrometer (Thermo Fisher Scientific). Peptides were separated on a reversed-phase column using a linear gradient of 3–28% ACN in 0.1% FA over 90 min, 28-40% ACN over 5 min and 40-80% ACN over 5 min at a flow rate of 300 nL/min. Full MS1 scans were collected over an m/z range of 350–950 at a resolution of 45,000, with an AGC target of 300% and a maximum injection time of 99.6 ms. Data-independent acquisition (DIA) was performed using 34 user-defined isolation windows at a resolution of 22,500 using higher-energy collisional dissociation (HCD) with a normalized collision energy of 27%, an AGC target of 3000%, and maximum injection time of 40.6 ms. Data was analysed with DIA-NN using an in-house spectral library, with up to two missed cleavages allowed, carbamidomethylation of cysteine set as a fixed modification, and methionine oxidation and protein N-terminal acetylation set as variable modifications. DIA-NN reports were filtered for high-confidence identifications with library, precursor, and protein group Q-values ≤ 0.01. Protein abundances were normalized using variance stabilization normalization^80^ and differential stability was analysed with R package limma using empirical Bayes moderation^81^. The proteins were considered significantly stabilized based on log2 fold change threshold 1.2 and adjusted p-value threshold 0.01.

### Molecular docking

The existing protein structures or structures of orthologous proteins (>30% identity) were downloaded from the PDB database^82^, otherwise the AlphaFold-predicted structures^83^ with removed low-confidence regions (<0.8) were used. The SMILES forms of compounds were converted to SDF format using Open Babel^84^. The charges were calculated using ADFR suite^85^ and the compounds were docked using Autodock Vina^86^. The figures were prepared with PyMoL (Schrödinger, LLC).

### Other statistical analyses

All statistical analyses were performed using basic R functions in R version 4.1.1 in RStudio (cite). The word clouds were created with the package wordcloud^87^, tables were handled and charts were plotted with tidyverse packages^88^. The gene enrichment was calculated by hypogeometric testing as described previously^89^. The figures were assembled with patchwork package ^90^ and Illustrator (Adobe Inc).

