## Extended Data 1 for "Thermal proteome profiling identifies new drug targets in Plasmodium falciparum parasites"

### Plasmepsin I (PF3D7_1407900)

Mefloquine

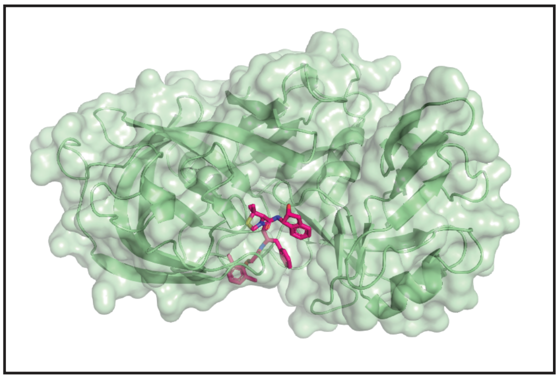

MMV006656

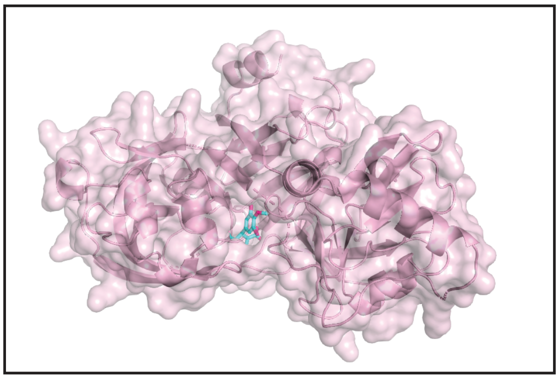

### Plasmepsin II (PF3D7_1408000)

Inhibitor

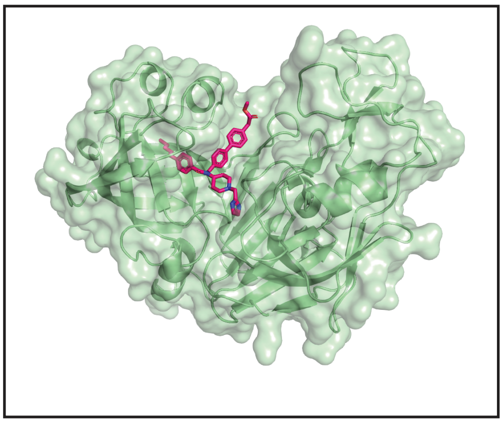

MMV007127

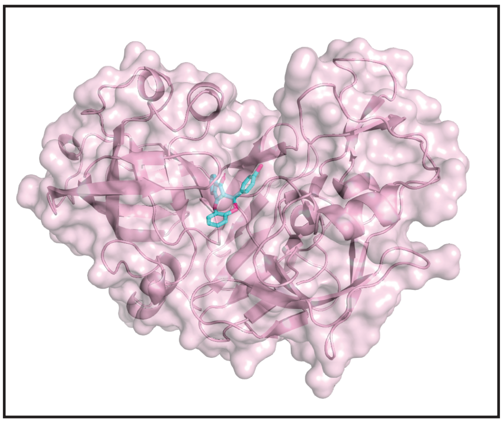

MMV020885

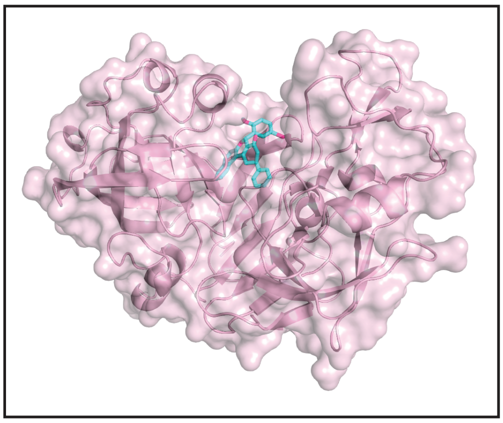

### Heat shock protein 90 (PF3D7_0708400)

ADP

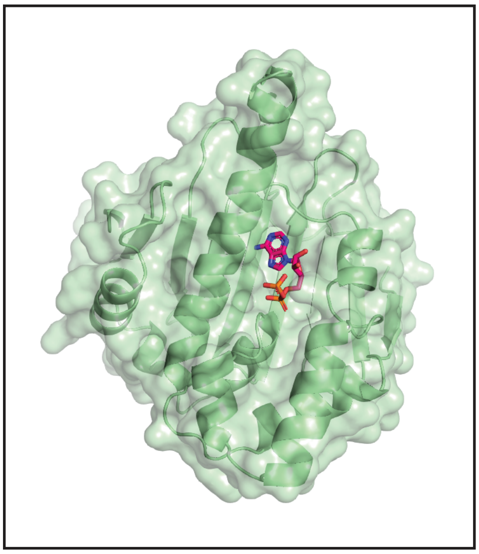

MMV066070

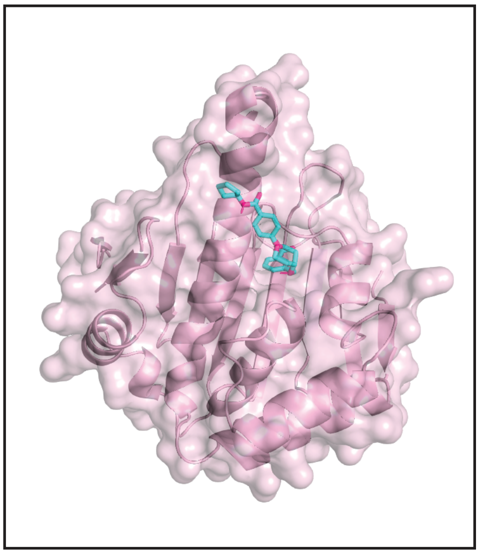

MMV665809

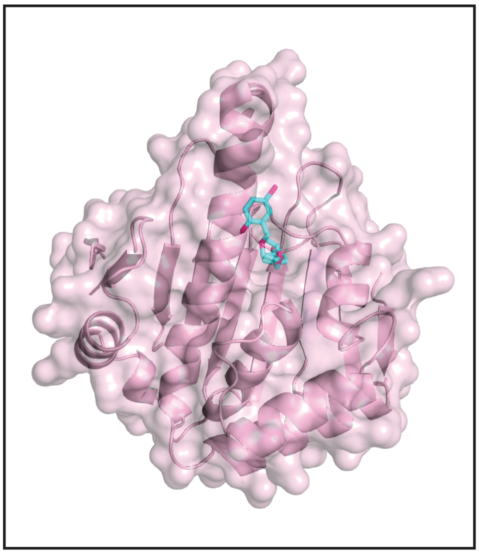

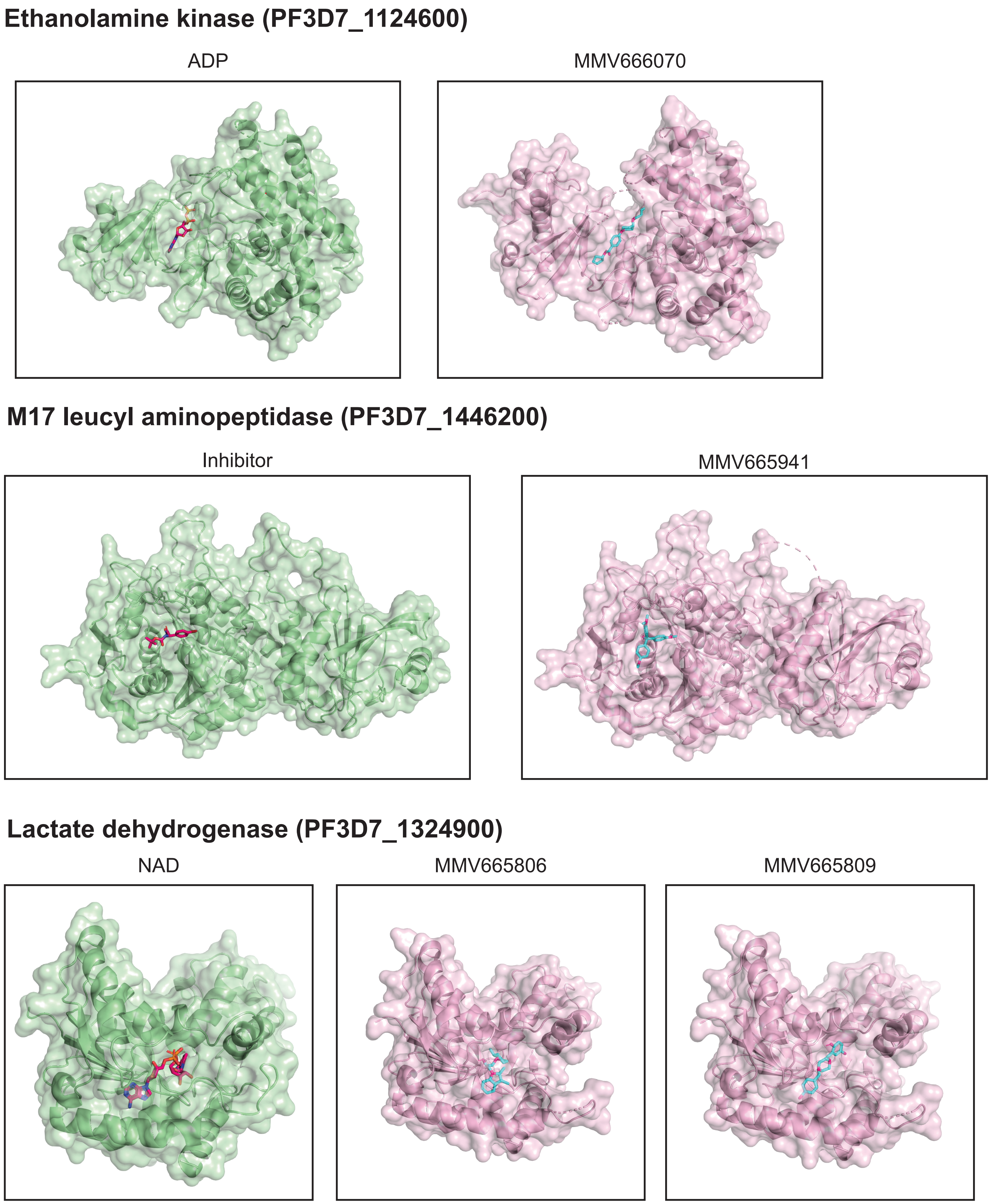
Ethanolamine kinase (PF3D7_1124600)
ADP
MMV666070
M17 leucyl aminopeptidase (PF3D7_1446200)
Inhibitor
MMV665941
Lactate dehydrogenase (PF3D7_1324900)
NAD
MMV665806
MMV665809

### Hexokinase (PF3D7_0624000)

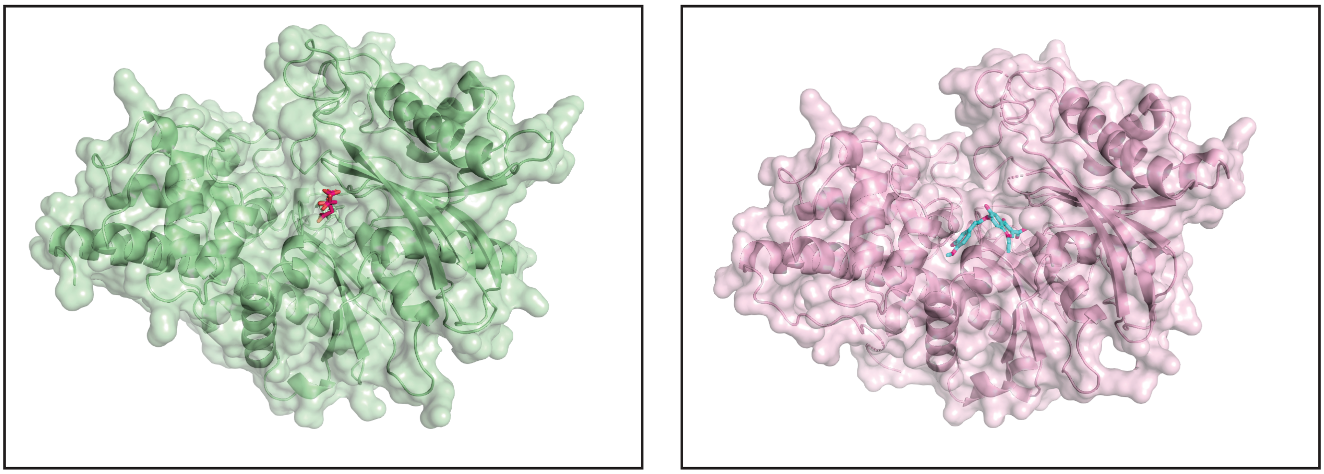


### Methionyl-tRNA synthetase (PF3D7_1034900)

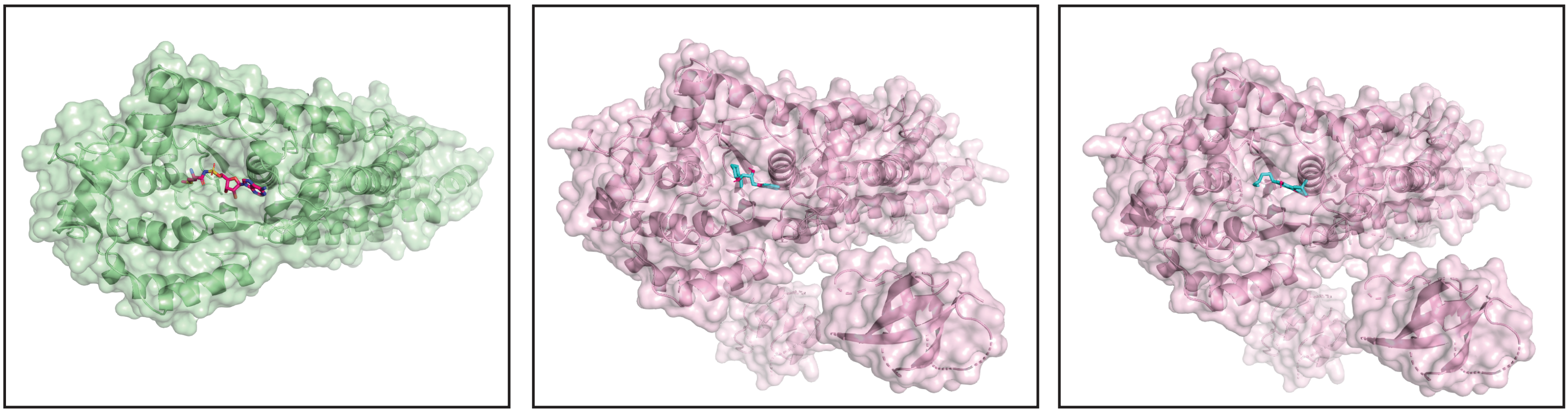


### Arginyl-tRNA synthetase (PF3D7_1218600)

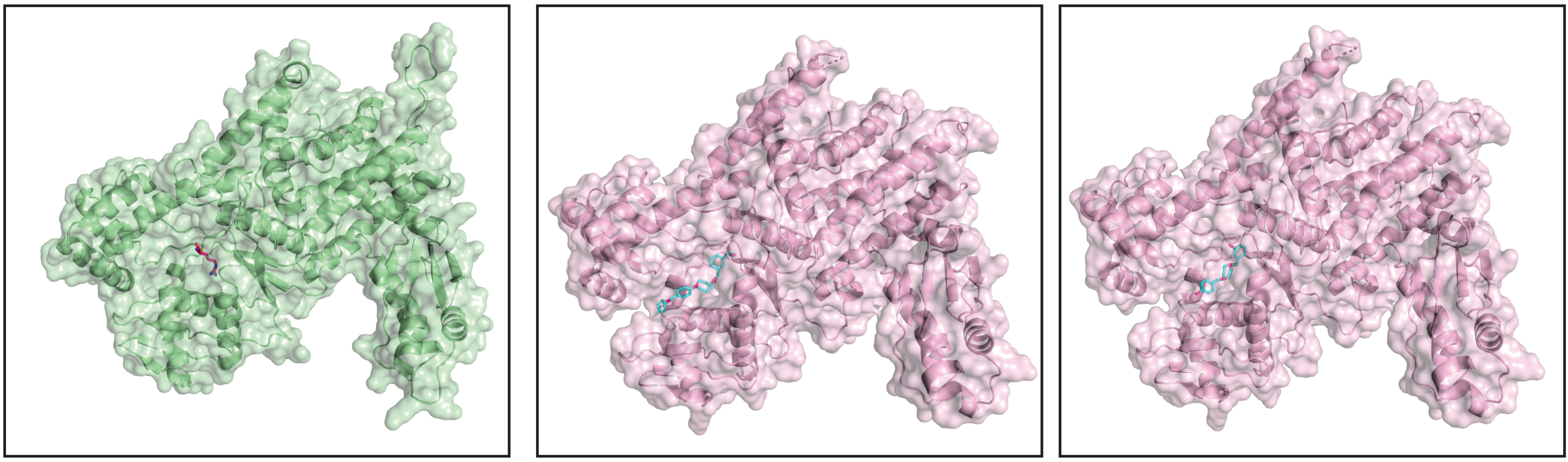

**Obg-like ATPase 1 (PF3D7_0722400)**

ATP analog

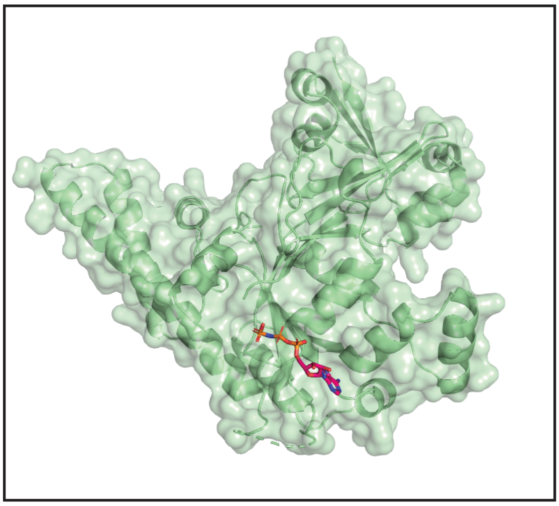

MMV665987

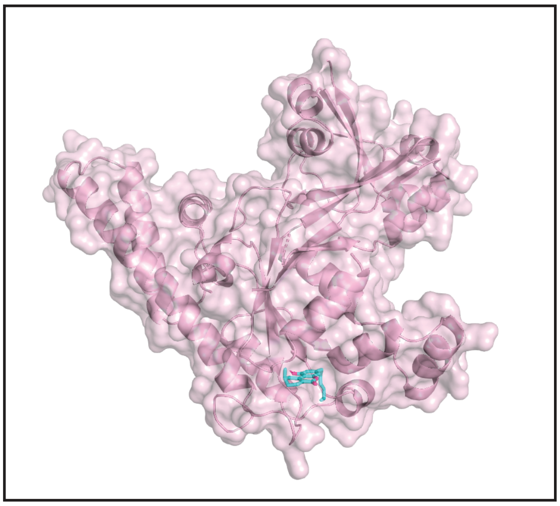

**Coproporphyrinogen-III oxidase (PF3D7_1142400)**

FICA

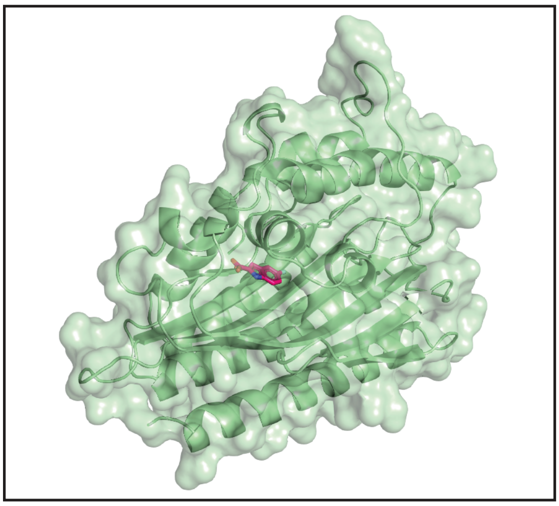

MMV006656

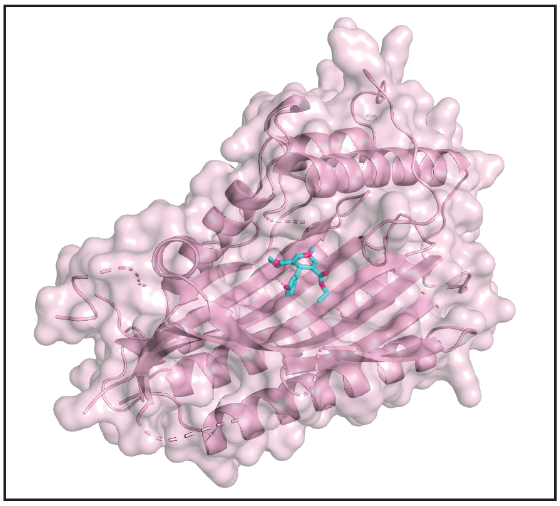

**Glutamyl-tRNA synthetase (PF3D7_1349200)**

ATP analog

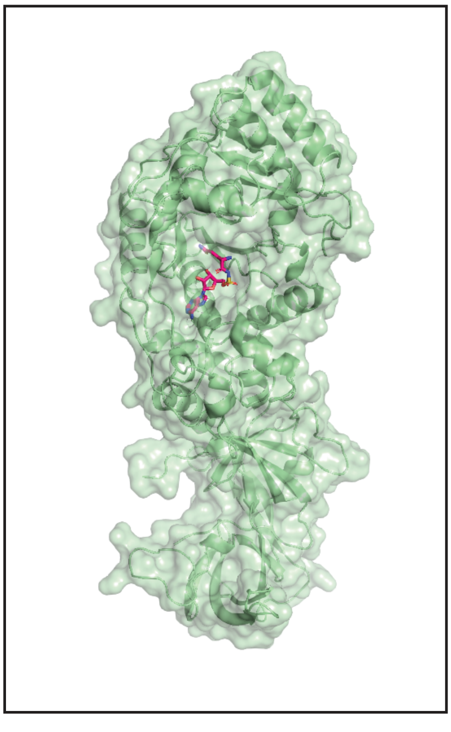

MMV007695

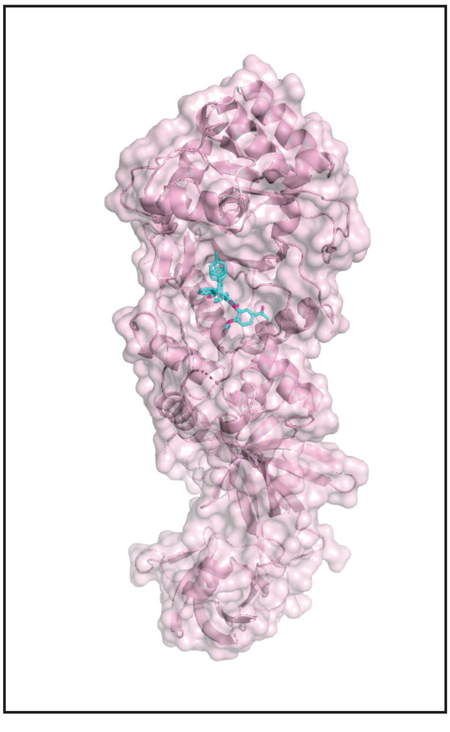
