## Supplementary Table 1 for "Thermal proteome profiling identifies new drug targets in Plasmodium falciparum parasites"

**Table 1.** 25 Screen compounds and information about their MoAs.

|  | **Prior information** | **Literature** |
| --- | --- | --- |
| MMV007127 | Male gametocyte-specific inhibitor. | ^1^ |
| MMV665915 | Inhibits lipid synthesis. | ^2^ |
| MMV665987 | No prior information. |  |
| MMV007695 | Metabolomic response similar to that of atovaquone. | ^3^ |
| MMV020885 | Causes defects in plastid segregation. | ^4^ |
| Cladosporin | Lysyl tRNA syntethase inhibitor. | ^5^ |
| MMV665971 | No prior information. |  |
| MMV006656 | Prevents rosette formation in blood group O red blood cells. | ^6^ |
|  | Moderate resistance conferred by PfATP4 mutation. | ^7^ |
|  | Putative PfATP4 inhibitor identified by Na+ uptake assay. | ^8^ |
| MMV665977 | Inhibits male gametocytogenesis. | ^1^ |
| GNF179 | Resistance conferred by mutations in PfAct and PfUGT. | ^9^ |
|  | Inhibits protein trafficking and establishment of permeation pathways, causes ER expansion | ^10^ |
| KAF156 (Ganaplacide) | Resistance conferred by mutations in PfAct and PfUGT. | ^9^ |
|  | Resistance conferred by mutations in PfCARL. | ^11^ |
|  | Inhibits protein trafficking and establishment of permeation pathways, causes ER expansion. | ^10^ |
| MMV396749 | Inhibits schizont egress. | ^4^ |
|  | Identified as PfATP4 inhibitor by Na+ uptake assay. | ^8^ |
| MMV1576856 | Racemic mixture of cipargamin and its enantiomer. Resistance to cipargamin is conferred by mutations in PfATP4 | ^12^ |
| NITD246 | Resistance conferred by mutations in PfATP4. | ^13^ |
| MMV665864 | No prior information. |  |
| MMV000848 | CETSA showed stabilization of PfPNP, crystal structure with PfPNP solved. | ^14^ |
|  | CETSA showed stabilization of falcilysin, crystal structure with falcilysin solved. | ^15^ |
| MMV665806 | CETSA showed stabilization of falcilysin, crystal structure with falcilysin solved. | ^15^ |
| MMV019127 | Inhibits schizont egress. | ^4^ |
| MMV665809 | No prior information. |  |
| MMV007574 | Fast-acting in asexual parasites. | ^16^ |
| MMV665886 | Inhibits translation in *Plasmodium falciparum.* | ^17^ |
| MMV665941 | Active against mature gametocytes. | ^18^ |
| MMV665794 | Although "irresistible", QRP1 mutations identified after exposure to mutator P. falciparum strain. | ^19^ |
|  | Overexpression of PfATP2 decreases susceptibility to this drug. PfATP2 knockdown induces hypersensitivity. Decreases cytosolic pH | ^20^ |
| MMV666070 | No prior information. |  |
| MMV665875 | Downregulates cysteine protease mRNA levels in *Theileria equi*. | ^21^ |
