## Supplementary Table 2 for "Thermal proteome profiling identifies new drug targets in Plasmodium falciparum parasites"

| 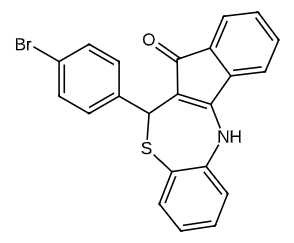**MMV007127** | 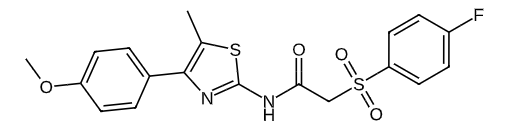**MMV665915** | 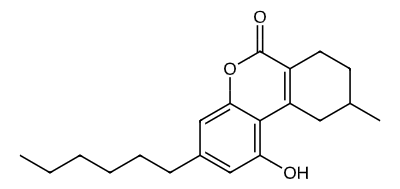**MMV665987** | **MMV007695**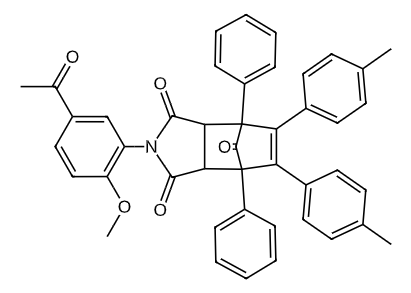 | 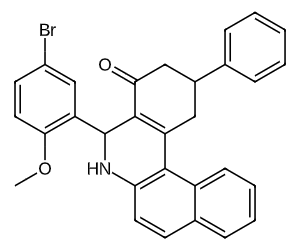**MMV020885** |
| --- | --- | --- | --- | --- |
| **Cladosporin**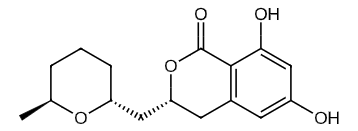 | **MMV665971**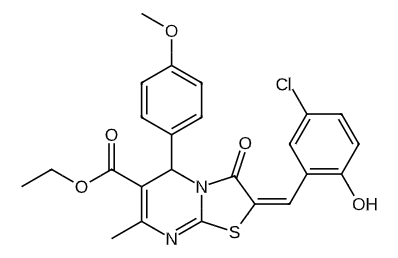 | 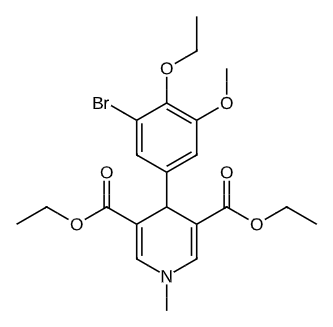**MMV006656** | **MMV665977**  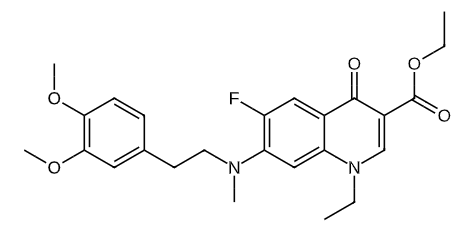 | 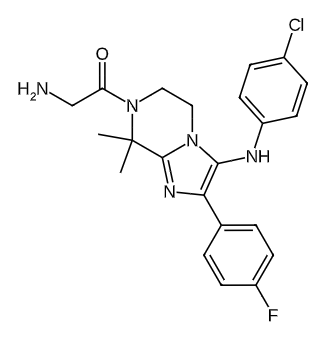**GNF179** |
| 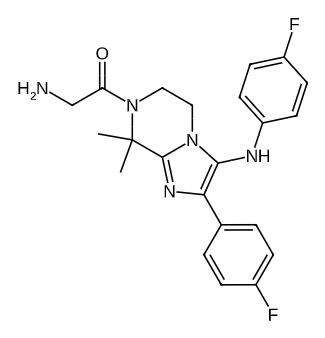**KAF156 (Ganaplacid)** | 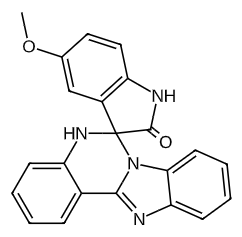**MMV396749** | **MMV1576856**  **(cipargamin enantiomer)**  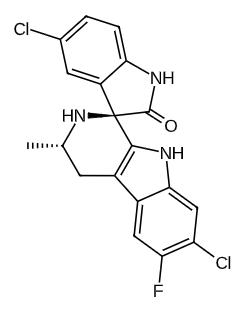 | **NIDT246**  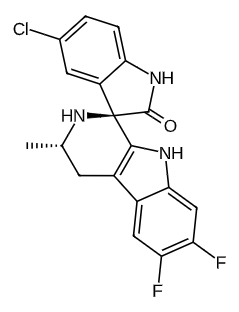 | **MMV665864**  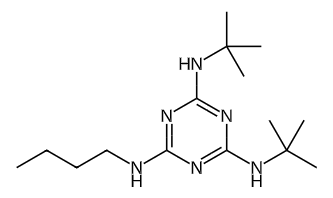 |
| **MMV000848**  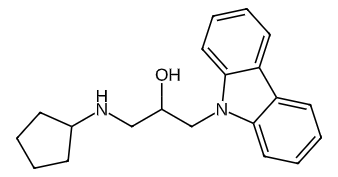 | **MMV665806**  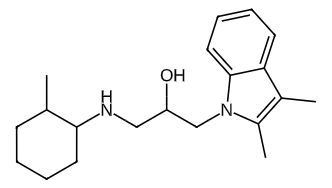 | **MMV019127**  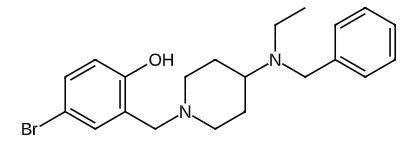 | **MMV665809**  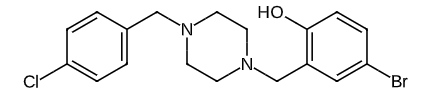 | **MMV007574**  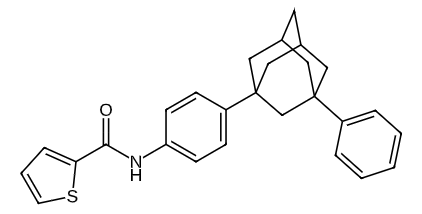 |
| **MMV665886**  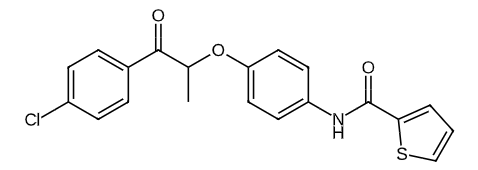 | **MMV665941**  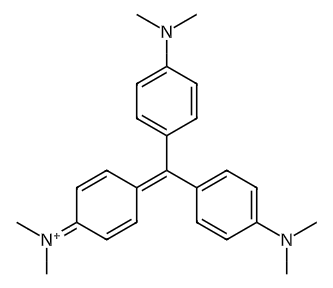 | **MMV665794**  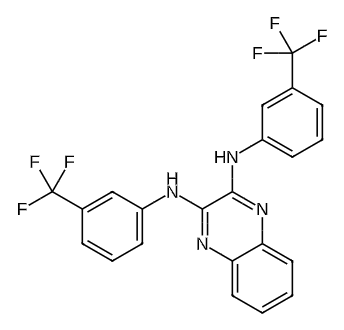 | **MMV666070**  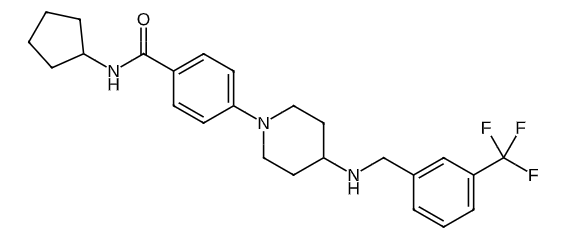 | **MMV665875**  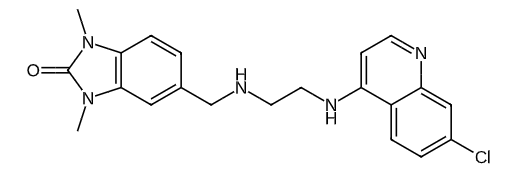 |
