## Supplementary Table 4 for "Thermal proteome profiling identifies new drug targets in Plasmodium falciparum parasites"

| Compound | Original BP ligand | Mean BE (kcal/mol) | No. PCs |
|---|---|---|---|
| **PF3D7_0513300 (purine nucleoside phosphorylase)** | | | |
| MMV000848 | MMV000848 | -8.143900 | 10 |
| MMV007695 | MMV000848 | -7.356286 | 7 |
| MMV066070 | MMV000848 | -9.221300 | 10 |
| **PF3D7_0624000 (hexokinase)** | | | |
| MMV665977 | Citrate | -6.475333 | 6 |
| **PF3D7_0708400 (heat shock protein 90)** | | | |
| MMV066070 | ADP | -7.995143 | 7 |
| MMV665809 | ADP | -6.459143 | 7 |
| **PF3D7_0722400 (Obg-like ATPase 1, putative)** | | | |
| MMV665987 | ATP analog | -6.933500 | 6 |
| **PF3D7_0907900 (peptide deformylase)** | | | |
| MMV665809 | PDF inhibitor | -7.360700 | 10 |
| MMV665971 | PDF inhibitor | -6.993500 | 10 |
| **PF3D7_1009000 (diphthine methyl ester synthase, putative)** | | | |
| MMV665915 | SAH | -8.272400 | 10 |
| MMV665977 | SAH | -7.635167 | 6 |
| **PF3D7_1034900 (methionine--tRNA ligase)** | | | |
| MMV665806 | ATP analog | -7.356375 | 8 |
| MMV665987 | ATP analog | -7.244667 | 9 |
| **PF3D7_1124600 (ethanolamine kinase)** | | | |
| MMV066070 | ADP | -8.890900 | 10 |
| **PF3D7_1142400 (coproporphyrinogen-III oxidase)** | | | |
| MMV006656 | FICA | -5.282714 | 7 |
| **PF3D7_1218600 (arginine--tRNA ligase)** | | | |
| MMV066070 | Arginine | -8.872556 | 9 |
| MMV665809 | Arginine | -6.956875 | 8 |
| **PF3D7_1324900 (L-lactate dehydrogenase)** | | | |
| MMV665806 | NAD | -6.913333 | 6 |
| MMV665809 | NAD | -6.756000 | 6 |
| **PF3D7_1343000 (phosphoethanolamine N-methyltransferase)** | | | |
| MMV066070 | AMQ (BP1) | -9.469200 | 5 |
| MMV066070 | AMQ (BP2) | -8.811200 | 5 |
| MMV066070 | SAM | -8.870200 | 5 |
| MMV665806 | AMQ (BP1) | -8.026300 | 10 |
| MMV665806 | AMQ (BP2) | -5.741778 | 9 |
| **PF3D7_1347200 (nucleoside transporter 1)** | | | |
| MMV020885 | GSK4 drug | -9.716600 | 5 |
| MMV066070 | GSK4 drug | -9.004200 | 10 |
| MMV665806 | GSK4 drug | -7.252333 | 9 |
| **PF3D7_1349200 (glutamate--tRNA ligase, putative)** | | | |
| MMV007695 | ATP analog | -7.773571 | 7 |
| **PF3D7_1350100 (lysine--tRNA ligase)** | | | |
| Cladosporine | Clado-like inhibitor | -7.019429 | 7 |
| Cladosporine | Lysine | -6.737857 | 7 |
| **PF3D7_1360800 (falcilysin)** | | | |
| MMV000848 | MFQ (BP1) | -7.643833 | 6 |
| MMV665806 | MFQ (BP1) | -7.345600 | 5 |
| **PF3D7_1407900 (plasmepsin I)** | | | |
| MMV006656 | KNI-10006 | -5.613250 | 8 |
| **PF3D7_1408000 (plasmepsin II)** | | | |
| MMV006656 | Inhibitor (BP1) | -5.582125 | 8 |
| MMV006656 | Inhibitor (BP2) | -5.549400 | 10 |
| MMV007127 | Inhibitor (BP1) | -8.129200 | 5 |
| MMV007127 | Inhibitor (BP2) | -8.070444 | 9 |
| MMV020885 | Inhibitor (BP1) | -8.840200 | 10 |
| MMV020885 | Inhibitor (BP2) | -8.838333 | 9 |
| **PF3D7_1446200 (M17 leucyl aminopeptidase)** | | | |
| MMV665941 | Inhibitor | -6.755778 | 9 |
